# Genome wide association study in 3,173 outbred rats identifies multiple loci for body weight, adiposity, and fasting glucose

**DOI:** 10.1101/422428

**Authors:** Apurva S. Chitre, Oksana Polesskaya, Katie Holl, Jianjun Gao, Riyan Cheng, Hannah Bimschleger, Angel Garcia Martinez, Tony George, Alexander F. Gileta, Wenyan Han, Aidan Horvath, Alesa Hughson, Keita Ishiwari, Christopher P. King, Alexander Lamparelli, Cassandra L. Versaggi, Connor Martin, Celine L. St. Pierre, Jordan A. Tripi, Tengfei Wang, Hao Chen, Shelly B. Flagel, Paul Meyer, Jerry Richards, Terry E. Robinson, Abraham A. Palmer, Leah C. Solberg Woods

## Abstract

**Objective:** Obesity is influenced by genetic and environmental factors. Despite success of human genome wide association studies (GWAS), the specific genes that confer obesity remain largely unknown. The objective of this study was to use outbred rats to identify genetic loci underlying obesity and related morphometric and metabolic traits.

**Methods:** We measured obesity-relevant traits including body weight, body length, body mass index, fasting glucose, and retroperitoneal, epididymal, and parametrial fat pad weight in 3,173 male and female adult N/NIH heterogeneous stock (HS) rats across three institutions, providing data for the largest rat GWAS to date. Genetic loci were identified using a linear mixed model that accounted for the complex family relationships of the HS and covariate to account for differences among the three phenotyping centers.

**Results:** We identified 32 independent loci, several of which contained only a single gene (e.g. *Epha5, Nrg1* and *Klhl14*) or obvious candidate genes (*Adcy3, Prlhr*). There were strong phenotypic and genetic correlations among obesity-related traits, and extensive pleiotropy at individual loci.

**Conclusions:** These studies demonstrate utility of HS rats for investigating the genetics of obesity related traits across institutions and identify several candidate genes for future functional testing.

## Introduction

Obesity is a growing health epidemic; over one third of the adult population and almost one fifth of all children in the United States are considered obese. There has been a steady increase in prevalence of obesity since the 1970’s, and prevalence is continuing to rise (1). Obesity is a major risk factor for multiple diseases including type 2 diabetes, cardiovascular disease, cancer, and stroke (2), thereby placing a tremendous burden on society. Obesity is caused by an interaction between genetic and environmental factors with genetic factors accounting for up to 70% of the population variance (3). Although human genome wide association studies (GWAS) of obesity have been extremely productive (4), much of the heritable variance is still unknown..

Model organism studies of body weight, morphometric, and metabolic traits represent a complementary approach to understanding the genetic basis of obesity. However, GWAS in model organisms are often limited by the modest recombination present in laboratory crosses. Heterogeneous stock (HS) rats were created by interbreeding eight inbred founder strains and were subsequently maintained as an outbred population (5), making them ideal for fine-mapping of genetic loci (6) Furthermore, HS founder strains have been fully sequenced (7), such that coding polymorphisms can be rapidly identified (8). Our group has previously mapped adiposity traits in HS rats using 743 male HS rats, which identified two genetic loci for visceral adiposity and body weight (8).

As part of a large, multi-site study (www.ratgenes.org), behavioral traits that are relevant to drug abuse have been assessed in thousands of male and female HS rats. To more fully utilize these rats, we have also collected phenotypic data on body weight and length (which permit calculation of body mass index (BMI)), fat pad weight and fasting glucose levels. All animals were also extensively genotyped. This dataset provides an unprecedented opportunity to map adiposity-related traits in a large cohort of HS rats.

## Materials and methods

### Animals

The NMcwi:HS colony (hereafter referred to as HS), was initiated by the NIH in 1984 using the following eight inbred founder strains: ACI/N, BN/SsN, BUF/N, F344/N, M520/N, MR/N, WKY/N and WN/N (5), which have previously been shown to differ significantly for adiposity traits (8). The rats described in this study were from generations 73 to 80, and were a separate cohort from previously published work (8). Breeders were given ad libitum access to Teklad 5010 diet (Envigo, Madison, Wisconsin).

The rats used for this study are part of a large multi-site project focused on genetic analysis of behavioral phenotypes related to drug abuse (www.ratgenes.org). HS rats from MCW were sent to three institutions throughout the United States: University of Tennessee Health Science Center (**TN**), University at Buffalo (**NY**), and University of Michigan (**MI**). Rats were shipped at 3-6 weeks of age and each site received multiple shipments over more than two years (from 10/27/2014 – 03/07/2017). There are multiple environmental differences between each site (described below), such that genetic loci will be independent from these environmental influences. Rats in TN were fed Teklad Irradiated LM-485 Mouse/Rat Diet (Envigo, Madison, Wisconsin); rats in NY were fed Teklad 18% Protein Rodent Diet (Envigo, Madison, Wisconsin), and rats in MI were fed Picolab Laboratory Rodent Diet Irradiated (LabDiet, St. Louis, Missouri).

Rats were exposed to a different battery of behavioral testing at each site (see **Supplementary Table S1)** followed by euthanasia, which occurred at different ages at each site. All phenotypes presented in this paper were collected at the time of euthanasia. Briefly, in MI, rats were housed in trios, exposed to a single modest dose of cocaine (15 mg/kg) each day for five days, and then euthanized 4-7 days after the final cocaine exposure (89 ± 6 days of age). In NY, rats were housed in pairs, tested for multiple behaviors over 16 weeks, exposed to a modest dose of cocaine (10 mg/kg) once daily for three days, and then euthanized 7-10 days after the last dose of cocaine (198 ± 13 days of age). In TN, there were two separate cohorts: breeders (sent from MCW) and experimental rats (bred in TN). Female breeders had mostly one, sometimes two litters and underwent no behavioral testing. The experimental rats were tested for multiple behaviors, exposed to nicotine (self-administration, resulting in a range of doses) for 12 days, and euthanized 10 days after the final dose of nicotine (73 ± 12 days of age). The number of rats phenotyped at each site, as well as ages when phenotypes were collected are shown in **Table 1**.

**Table 1.** Age and number of rats at time adiposity phenotypes were collected.

| Phenotyping site | Males, N | Females, N | Total N | Age, days, |
| --- | --- | --- | --- | --- |
| | | | | mean ±SD |
| MI | 519 | 514 | **1,033** | 89 +/- 6 |
| NY | 449 | 443 | **892** | 198 +/- 13 |
| TN (experimental) | 458 | 451 | **909** | 73 +/- 12 |
| TN (breeders) | 182 | 157 | **339** | 169 +/- 34 |
|  | 1,608 | 1,565 | **3,173** |  |

### Phenotyping and Tissue collection

Several days after completion of behavioral experiments (see above), rats were fasted overnight (17 +/-2 hours), and body weight was measured. Under anesthesia (phentobarbital at MI and NY, isoflurane at TN), two measures of body length (from nose to base of the tail (body length_NoTail) and from nose to end of tail (body length_Tail)) were collected, allowing us to calculate tail length (TL) and two measures of body mass index: BMI_NoTail and BMI_Tail. BMI was calculated as: (body weight/body length^2^) x 10. For animals in MI and NY, we also measured fasting glucose levels using the Contour Next EZ system (Bayer, Elkhart, IN). Several tissues were dissected and weighed including retroperitoneal (RetroFat), epididymal (EpiFat; males), and parametrial (ParaFat; females) visceral fat pads. All protocols were approved by the Institutional Animal Care and Use Committees (**IACUC**) for each of the relevant institutions.

### Genotyping

Genotypes were determined using genotyping-by-sequencing (**GBS**), as described previously(9, 10).. This produced 3,400,759 SNPs with an estimated error rate <1%. Variants for X- and Y-chromosomes were not called. Prior to GWAS, SNPs in high linkage disequilibrium were removed using PLINK (11) with r^2^ cut-off 0.95; this produced set of 128,447 SNPs which were used for GWAS, genetic correlations, and heritability estimates. The unpruned set of SNPs was used to produce LocusZoom plots (12).

### Phenotypic and genetic correlations and heritability estimates

Each trait within each research site was quantile normalized separately for males and females, this approach is similar to using sex as a covariate. Other relevant covariates (including age, batch number and dissector) were identified for each trait, and covariate effects were regressed out if they were significant and explained more than 2% of the variance (see **Supplementary Table S2**). Residuals were then quantile normalized again, after which the data for each sex and site was pooled prior to further analysis. This approach removed mean differences due to sex, however it did not attempt to model gene by sex interactions. By quantile normalizing the three centers separately, we also addressed the numerous and confounded differences between the three cohorts, such that QTL identified in the current study are resistant to environmental influences between the sites. (see **Supplementary Table S1**) Phenotypic correlations were determined using Spearman’s test. Genetic correlations were calculated using bivariate GREML analysis as implemented by GTCA(13, 14). GCTA-GREML analysis was used to estimate proportion of variance attributable to SNPs.

### Genetic mapping

GWAS analysis employed a linear mixed model, as implemented in the software GEMMA(15), using dosages and genetic relatedness matrices (GRM) to account for the complex family relationships within the HS population and the Leave One Chromosome Out (LOCO) method to avoid proximal contamination (16, 17). Significance thresholds were calculated using permutation, because all traits were quantile normalized, we used the same threshold for all traits (18). To identify QTLs, we scanned each chromosome to determine if there was at least one SNP that exceeded the permutation-derived threshold of –log_10_(p) > 5.6, that was supported by a second SNP within 0.5 Mb that had a p-value that was within 2 – log_10_(p) units of the index SNP. This algorithm failed to identify 2 SNPs, which were then identified by using wider supporting interval (chr10:85082795 for body length_tail and chr12:5782829 for RetroFat). Then correlation between the top SNP and other SNPs in the 6 Mb vicinity was calculated. Other QTLs on the same chromosome were tested to ensure that they were independent of the first. To establish independence, we used the top SNP from the first QTL as a covariate and performed a second GWAS. If the resulting GWAS had an additional SNP (on the same chromosome) with a p-value that exceeded our permutation-derived threshold, it was considered to be a second, independent locus. This process was repeated (including all previously significant SNPs as covariates), until no more QTLs were detected on a given chromosome.

Linkage disequilibrium (LD) intervals for the identified QTL were determined by identifying those markers that had a high correlation coefficient with the peak marker (r^2^ = 0.6). Credible set analysis (19) was also performed for each locus. The credible set analysis uses a Bayesian approach to calculate the posterior probability for each SNP (“probability of being causal”). This method also chooses the credible set of SNPs, that is the smallest set of SNPs that can account for 99% of the posterior probability.

## Results

### Strong phenotypic and genetic correlations between multiple adiposity traits

We observed strong phenotypic and genetic correlations between adiposity traits (**Figure 1**). Although the phenotypic correlation was weaker, fasting glucose levels were correlated with body weight, BMI_Tail, BMI_NoTail, RetroFat and EpiFat, but not with body length or ParaFat. Genetic correlations for fasting glucose levels differ from the phenotypic correlations, with a negative correlation seen with body length_NoTail and a positive correlation with ParaFat and BMI_NoTail. TL exhibits strong phenotypic and genetic correlations with BMI_NoTail, and negative phenotypic and non-existent genetic correlations with BMI_Tail. (**Supplementary Figure S1**) Tail length is also moderately correlated with the adiposity traits, indicating a role for this trait in rat metabolism, possibly through energy expenditure as rats lose heat through their tail.

**Figure 1.**
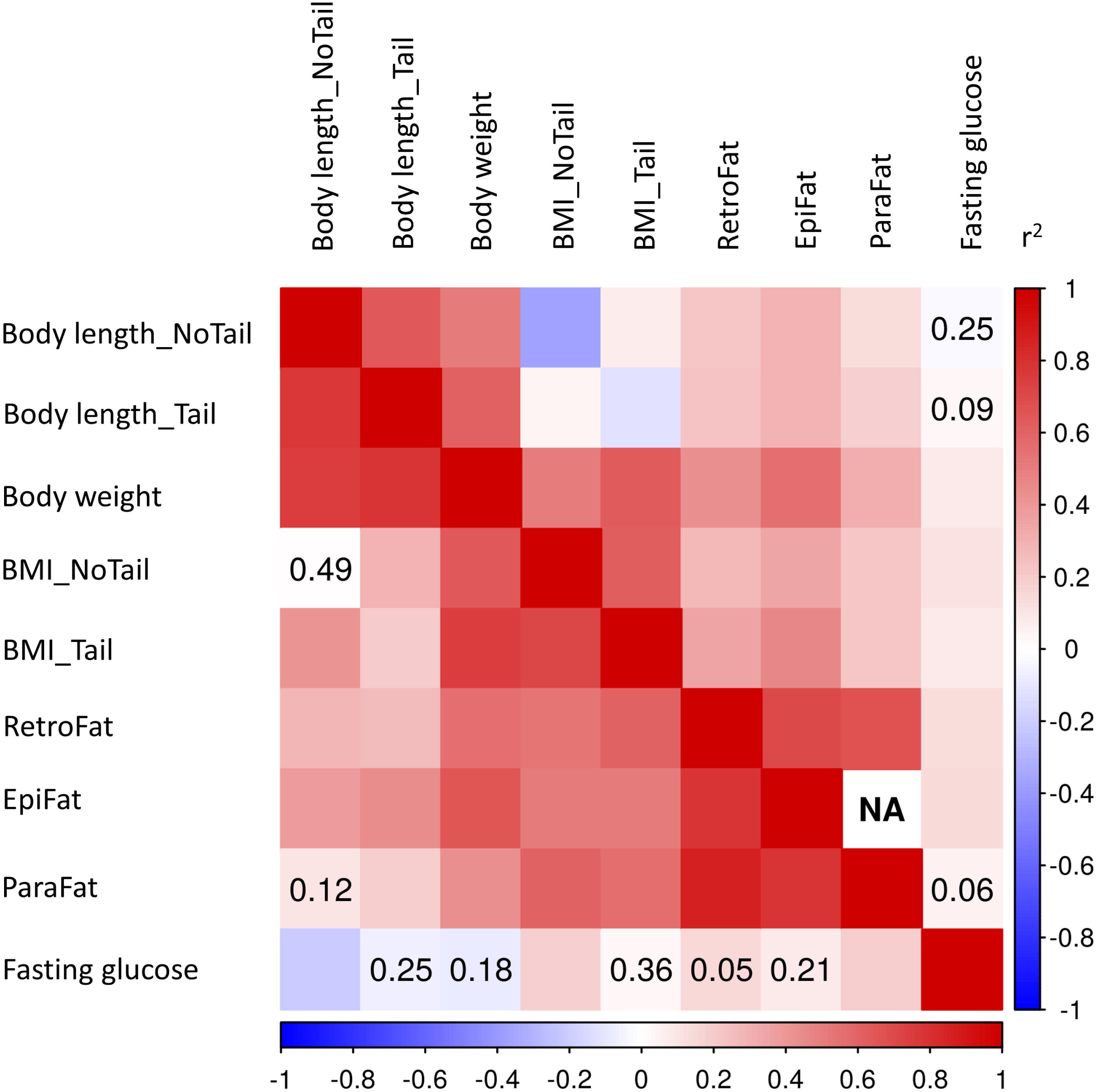
Genetic and phenotypic correlation between adiposity traits and fasting glucose. Phenotypic correlations are depicted in the upper part, genetic – in the lower part of the matrix. Number inside squares show p-value > 0.05.

### Adiposity traits exhibit high heritability

Heritability estimates for adiposity traits ranged from 0.26 ± 0.03 (BMI_NoTail) to 0.46 ± 0.03 (body weight; **Table 2**), while heritability estimates for fasting glucose were lower. We also calculated heritability for males and females separately, in general the results were similar for the two sexes (**Supplementary Table S3)**. Although we performed the GWAS separately for both sexes, we have not provided the results of sex specific GWAS because the reduction in power dramatically reduced the number of genome wide significant results.

**Table 2.**
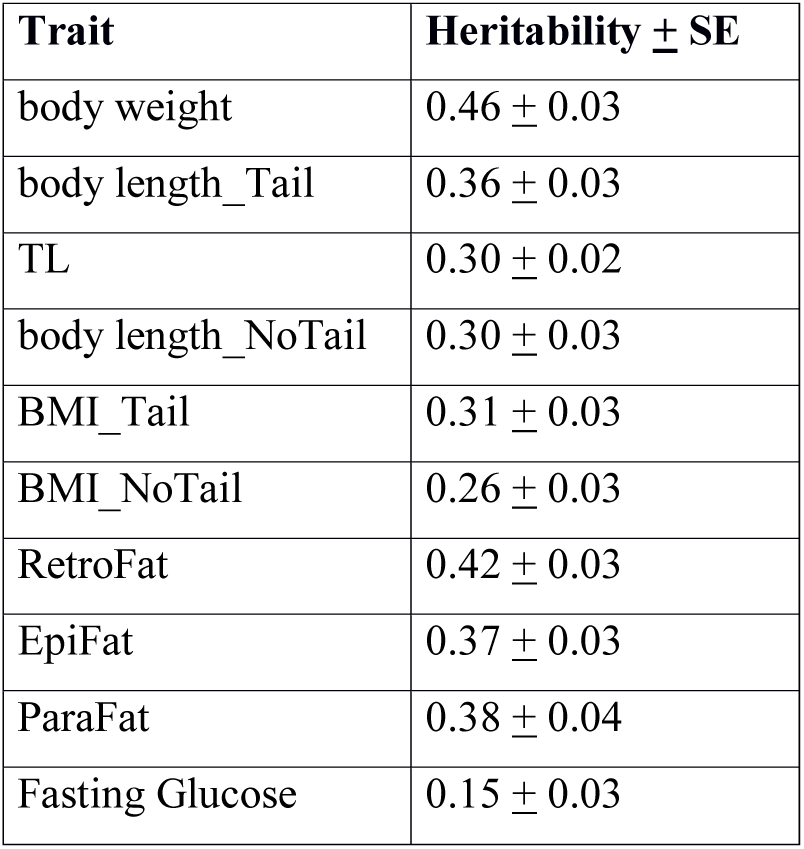
SNP Heritability estimates.

| Trait | Heritability ± SE |
| --- | --- |
| body weight | 0.46 ± 0.03 |
| body length_Tail | 0.36 ± 0.03 |
| TL | 0.30 ± 0.02 |
| body length_NoTail | 0.30 ± 0.03 |
| BMI_Tail | 0.31 ± 0.03 |
| BMI_NoTail | 0.26 ± 0.03 |
| RetroFat | 0.42 ± 0.03 |
| EpiFat | 0.37 ± 0.03 |
| ParaFat | 0.38 ± 0.04 |
| Fasting Glucose | 0.15 ± 0.03 |

### Identification of multiple GWAS hits

We identified a total of 32 independent loci for eight adiposity traits (**Figure 2**). Six of these loci map to more than one trait (**Figure 2)**, four of which are likely pleiotropic (see below and **Table 3**). Specifically, we identified nine QTLs for body weight, seven for RetroFat, three for EpiFat, one for ParaFat, eight for Body length_Tail, five for Body length_NoTail, three for BMI_Tail and three for BMI_No Tail. The most highly significant loci are pleiotropic: chr. 1:282 (Body weight, BMI_Tail, RetroFat, EpiFat, ParaFat), chr6: 27 Mb (RetroFat, EpiFat) and chr. 7: 36Mb (Body weight, Body length_NoTail, Body length_Tail, TL). Body weight QTL consistently overlaps with both fat pad QTL and Body length QTL, while only one BMI_Tail QTL overlaps with adiposity traits. BMI_Tail and BMI_NoTail loci do not overlap. It is difficult to use these data to determine if the BMI calculation is useful, although the fact that BMI_Tail maps to the highly significant pleiotropic locus on chr. 1: 282 Mb suggests similar biology may underlie these traits. We identified two loci for fasting glucose, one of which overlaps a Body weight locus on chromosome 10, although it is unlikely that these loci are driven by the same variant (see below). Four QTL are identified for TL, one of which overlaps with previously identified loci (see **Supplementary Figure S2**). LD intervals were 0.2-9.2 Mb and contained anywhere from 1 to 96 genes. **Table 3** provides a detailed summary of those findings; **Supplementary Table S3** contains all the information from **Table 3** with additional information such as the strain distribution pattern (SDP) of the founder strains as well as the full list of genes within each interval, and credible set analysis results. Manhattan plots for all traits are included in **Supplementary Figure S2**.

**Table 3.** Summary of QTLs.

| Trait | Peak Marker |  |  |  |  | LD Interval |  |  | Genes in the LD interval |
| --- | --- | --- | --- | --- | --- | --- | --- | --- | --- |
|  | Position | -logP | Ref Allele | Allele frequency | Effect size | start (bp) | stop (bp) | size (Mb) |  |
| BMI_Tail | chr1:106866154 | 6.05 | G | 0.29 | 0.15 ± 0.03 | 105,730,059 | 109,396,142 | 3.67 | 7 |
| RetroFat | chr1:160530456 | 7.51 | T | 0.53 | -0.14 + 0.02 | 157,254,290 | 162,857,247 | 5.60 | 15 |
| Body Weight | chr1:185730317 | 7.58 | C | 0.74 | 0.16 ± 0.02 | 184,463,432 | 187,738,111 | 3.27 | 6 |
| BMI_NoTail | chr1:187300775 | 6.72 | A | 0.66 | 0.15 ± 0.02 | 184,772,656 | 189,346,447 | 4.57 | 27 |
| TL | chr1:253524003 | 5.76 | G | 0.45 | 0.13 ± 0.03 | 253,082,171 | 254,725,734 | 1.64 | 6 |
| ParaFat | chr1:280924549 | 7.22 | G | 0.57 | 0.20 ± 0.03 | 280,876,316 | 282,114,080 | 1.24 | 9 |
| Body Weight | chr1:281756885 | 16.21 | C | 0.58 | 0.21 ± 0.02 | 280,924,333 | 282,114,080 | 1.19 | *Fam204a, Prlhr, Cacul1* |
| RetroFat | chr1:281777218 | 20.21 | A | 0.57 | 0.24 + 0.02 | 280,924,333 | 282,114,080 | 1.19 | 9 |
| EpiFat | chr1:281802657 | 14.00 | C | 0.56 | 0.29 ± 0.03 | 280,924,333 | 282,736,277 | 1.81 | 13 |
| BMI_Tail | chr1:282049439 | 11.44 | C | 0.55 | 0.18 ± 0.02 | 280,924,333 | 282,736,277 | 1.81 | 13 |
| Body Weight | chr2:65816485 | 6.55 | A | 0.91 | 0.20 ± 0.03 | 62,570,942 | 71,814,490 | 9.24 | 6 |
| RetroFat | chr3:95389621 | 13.12 | A | 0.44 | -0.19 + 0.02 | 92,336,188 | 97,685,154 | 5.35 | 30 |
| Body Length_NoTail | chr3:136021511 | 6.92 | A | 0.77 | -0.16 ± 0.03 | 132,291,573 | 137,146,532 | 4.85 | 10 |
| Body Length_Tail | chr3:136021511 | 5.97 | A | 0.77 | -0.14 ± 0.03 | 132,291,573 | 137,146,532 | 4.85 | 10 |
| Body Weight | chr3:136021511 | 7.34 | A | 0.77 | -0.16 ± 0.02 | 132,291,573 | 137,146,532 | 4.85 | 10 |
| RetroFat | chr3:137537161 | 8.21 | G | 0.44 | -0.15 + 0.02 | 136,161,761 | 138,849,437 | 2.69 | 20 |
| Body Weight | chr5:50933779 | 6.99 | A | 0.78 | -0.16 ± 0.03 | 49,152,709 | 50,940,275 | 1.79 | 12 |
| EpiFat | chr6:26266960 | 7.25 | G | 0.59 | -0.20 ± 0.03 | 22,684,886 | 28,223,800 | 5.54 | 55 |
| RetroFat | chr6:28148338 | 19.50 | G | 0.65 | -0.26 + 0.02 | 25,954,450 | 28,752,109 | 2.80 | 58 |
| Body Length_Tail | chr6:137745191 | 5.89 | C | 0.54 | -0.12 ± 0.02 | 136,769,305 | 138,087,183 | 1.32 | 24 |
| Body Weight | chr7:24886476 | 7.29 | G | 0.07 | 0.26 ± 0.04 | 24,869,890 | 25,213,445 | 0.34 | *Tcp1l12, Nuak1, Ckap4* |
| Body Weight | chr7:36497588 | 9.77 | C | 0.14 | -0.24 ± 0.03 | 34,119,928 | 36,522,260 | 2.40 | 21 |
| Body Length_NoTail | chr7:36517726 | 7.88 | T | 0.12 | -0.25 ± 0.04 | 34,119,928 | 36,522,260 | 2.40 | 15 |
| Body Length_Tail | chr7:36517726 | 14.37 | T | 0.12 | -0.33 ± 0.04 | 34,119,928 | 36,522,260 | 2.40 | 21 |
| TL | chr7:36526715 | 6.10 | A | 0.37 | 0.14 ± 0.03 | 36,289,478 | 36,765,903 | 0.48 | 5 |
| Body Length_Tail | chr8:118711320 | 5.72 | A | 0.67 | 0.13 ± 0.02 | 116,614,891 | 119,781,444 | 3.17 | 96 |
| Body Length_Tail | chr9:65078205 | 6.81 | C | 0.76 | 0.16 ± 0.03 | 64,013,585 | 65,670,399 | 1.66 | 20 |
| BMI_Tail | chr10:84080794 | 8.21 | G | 0.57 | -0.16 ± 0.02 | 83,442,285 | 85,006,252 | 1.56 | 42 |
| TL | chr10:84263936 | 9.38 | T | 0.57 | 0.18 ± 0.03 | 83,442,285 | 85,006,252 | 1.56 | 42 |
| Body Length_Tail | chr10:85082795 | 7.37 | C | 0.72 | 0.17 ± 0.03 | 84,902,901 | 85,239,899 | 0.34 | 10 |
| BMI_NoTail | chr10:96804258 | 7.66 | T | 0.48 | -0.16 ± 0.02 | 96,561,667 | 98,097,621 | 1.54 | 16 |
| Fasting Glucose | chr10:109944213 | 6.33 | A | 0.75 | -0.18 ± 0.03 | 108,350,175 | 110,315,359 | 1.97 | 49 |
| Body Weight | chr10:111010289 | 6.10 | T | 0.41 | 0.13 ± 0.02 | 110,664,101 | 111,022,246 | 0.36 | *Tbcd, Znf750, B3gntl1, Metrnl* |
| Body Length_Tail | chr12:2199384 | 8.94 | C | 0.18 | -0.22 ± 0.03 | 846,435 | 5,587,624 | 4.74 | 50 |
| Body Weight | chr12:5738696 | 9.12 | C | 0.59 | 0.16 ± 0.02 | 4,938,015 | 6,078,451 | 1.14 | 6 |
| RetroFat | chr12:5782829 | 7.12 | T | 0.73 | 0.14 + 0.02 | 455,837 | 6,259,634 | 5.80 | 59 |
| Body Length_NoTail | chr12:6239515 | 6.26 | A | 0.52 | 0.13 ± 0.02 | 6,078,237 | 7,035,028 | 0.96 | 9 |
| Body Length_Tail | chr12:43060205 | 6.27 | G | 0.75 | -0.15 ± 0.02 | 42,024,283 | 43,190,584 | 1.17 | *Oas1g, Tbx3, Tbx5* |
| TL | chr13:109566014 | 7.04 | T | 0.34 | 0.15 + 0.03 | 108,055,241 | 111,868,706 | 3.81 | 30 |
| RetroFat | chr13:55021887 | 7.16 | C | 0.67 | 0.14 + 0.02 | 54,910,667 | 56,487,438 | 1.58 | 7 |
| Body Length_NoTail | chr14:26365986 | 6.54 | A | 0.73 | -0.16 ± 0.03 | 25,129,356 | 26,478,660 | 1.35 | *Epha5* |
| Fasting Glucose | chr14:86029588 | 6.27 | T | 0.27 | -0.17 ± 0.03 | 82,473,632 | 87,575,807 | 5.10 | 34 |
| Body Length_NoTail | chr16:64014119 | 6.35 | G | 0.55 | 0.13 ± 0.02 | 64,003,027 | 64,416,540 | 0.41 | *Nrg1* |
| EpiFat | chr18:12674871 | 6.47 | G | 0.22 | -0.22 ± 0.04 | 12,597,378 | 12,792,162 | 0.19 | *Klhl14* |
| BMI_NoTail | chr18:25190274 | 5.78 | T | 0.04 | -0.34 ± 0.07 | 23,785,532 | 25,418,376 | 1.63 | 20 |
| TL | chr19:32004685 | 7.81 | C | 0.41 | -0.15 ± 0.03 | 31,176,512 | 32,698,569 | 1.52 | 7 |

**Figure 2.**
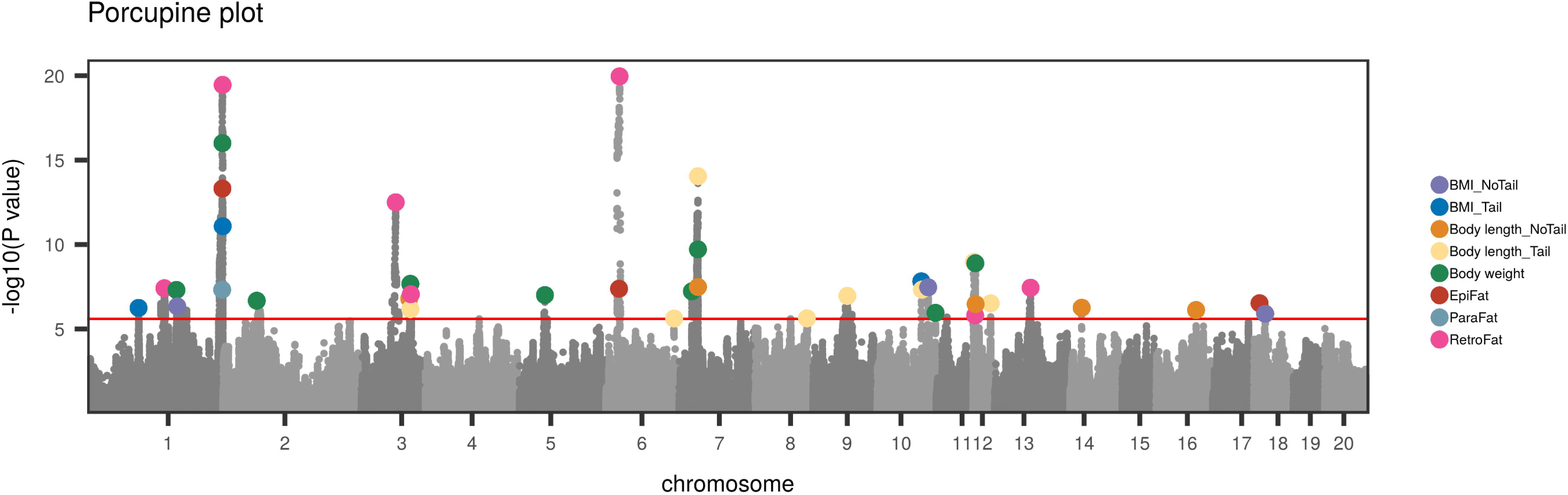
Combined Manhattan plots of GWAS data for eight adipose traits. Genome-wide association results from the GWA analysis. The chromosomal distribution of all the P-values (- log10 P-values) is shown, with top SNPs highlighted.

### Pleiotropic Loci

To determine if traits that mapped to the same location are pleiotropic, we considered the mean allele frequency (**MAF**), and the SDP of the index SNP among the 8 founder strains that were used to create the HS. In a few cases, we found that loci for two or more phenotypes overlapped in terms of their chromosomal positions, but had different MAF and SDP, suggesting different causal loci that happened to be located in approximately the same genetic location. We also observed a few cases where the SDP was similar but not identical (e.g. one strains’ genotype was dissimilar between the two loci), we assumed that those situations were consistent with pleiotropy. Note that this approach is conceptually similar to the estimation of the phenotype associated with founder haplotypes which was described in Svenson et al.(20) and in Yalcin et al. (21) but would not perform well if there were more than two causal alleles.

Using these criteria, we designated four loci as pleiotropic, which are highlighted in purple in **Table 3**. The chr 1: 281 Mb is the strongest, with –log_10_P values that range from 7-20 for nearly all of the adiposity traits, including body weight, BMI_Tail, and all three fat pads (RetroFat, EpiFat and ParaFat). Although BMI_Tail maps to the pleitropic locus on chr 1: 281 Mb, BMI_NoTail does not.. In addition chr 6: ∼27 Mb maps both RetroFat and EpiFat and Finally, chr 3 and 7 map body weight, body length_Tail and body length_NoTail. Nearby loci that were not considered pleotropic are highlighted in grey in **Table 3**. We identified two sets of QTLs that mapped to similar regions, and therefore might have appeared to be pleiotropic; however, after applying the criteria described above, we determined that they were not truly pleiotropic because the MAF and SDP of the founder strains (see **Supplementary Table S3**) were not similar. These loci were chr 10: ∼110 Mb for fasting glucose and body weight and chr 12: 6 Mb for body weight, RetroFat and body length_NoTail.

### Candidate gene identification

The number of genes within the identified QTL ranges from 1 to 96 (**Table 3**). There are three regions that contain a single gene: *Epha5* within chr 14: 26 Mb for body length_NoTail, *Nrg1* within chr 16: 64 Mb for body length_NoTail and *Klh14 within* chr18: 12 Mb for EpiFat. Both *Epha5* and *Nrg1* are known to be involved in growth and development and are therefore logical candidate genes. In contrast, very little is known about *Klhl14*, meaning that if this gene is responsible for the observed QTL it would offer novel biological insight. Despite the fact that these loci only contained a single gene, it is possible that the causal allele is a regulatory variant that is located in this interval but regulates a gene outside of the identified interval.

All other loci contained more than one gene. To identify candidate genes, we used founder sequence information (22) to identify potentially damaging coding variants using the software SnpEff. (23) We identified 152 coding variants for 18 QTL, two of which were predicted to have high impact. The SDP for 63 coding variants matched the SDP at the peak SNP, and are therefore candidates for future analysis. (**Supplementary Table S4**) Combining these data with a literature search, we identified plausible candidate genes/variants within six loci. (**Table 4**) Representative LocusZoom plots for select loci are shown in **Figure 3**, locus zoom plots for all other loci are in **Supplementary Figure S3**.

**Table 4.** Moderate/High impact coding variants that fall within adiposity QTL and are supported by SDP and/or literature search.

| Trait | QTL | Candidate Gene | SNP<br>In LD<br>With<br>Peak<br>Marker | SNP.<br>change | Amino.<br>acid.<br>change | effect | impact | ACI | BN | BUF | F344 | M520 | MR | WN | WKY | Literature Support |
| --- | --- | --- | --- | --- | --- | --- | --- | --- | --- | --- | --- | --- | --- | --- | --- | --- |
| Fasting Glucose | chr10:<br>109 Mb | *Rnf213* | chr10:108582280 | c.9502C>G | p.His3168Asp | missense_variant | MODERATE | GG | CC | GG | GG | GG | GG | GG | GG | PMID 2341075 |
|  |  |  | chr10:108582544 | c.9766T>C | p.Trp3256Arg | missense_variant | MODERATE | CC | TT | CC | CC | CC | CC | CC | CC |  |
|  |  |  | chr10:108604073 | c.12263G>C | p.Ser4088Thr | missense_variant | MODERATE | CC | GG | CC | CC | CC | CC | CC | CC |  |
| Fasting Glucose | chr14:<br>86 Mb | *Xbp1* | chr14:85756355 | c.397A>G | p.Asn133Asp | missense_variant | MODERATE | AA | AA | AA | AA | AA | AA | AA | GG | 366 Pubmed articles for Xbp1, glucose |
| BMI_Tail | chr1:<br>282 Mb | *Prlhr* | chr1:281755911 | c.3G>A | p.Met1? | start_lost | HIGH | CC | CC | TT | CC | CC | CC | CC | TT | PMID 29193816 |
| Body Weight |  |  | chr1:281755911 | c.3G>A | p.Met1? | start_lost | HIGH | CC | CC | TT | CC | CC | CC | CC | TT |  |
| ParaFat |  |  | chr1:281755911 | c.3G>A | p.Met1? | start_lost | HIGH | CC | CC | TT | CC | CC | CC | CC | TT |  |
| RetroFat |  |  | chr1:281755911 | c.3G>A | p.Met1? | start_lost | HIGH | CC | CC | TT | CC | CC | CC | CC | TT |  |
| EpiFat |  |  | chr1:281755911 | c.3G>A | p.Met1? | start_lost | HIGH | CC | CC | TT | CC | CC | CC | CC | TT |  |
| RetroFat | chr6:<br>28 Mb | *Adcy3* | chr6:28572363 | c.362T>C | p.Leu121Pro | missense_variant | MODERATE | TT | TT | TT | TT | TT | TT | TT | CC | PMID 29193816 |
| Body Length_Tail | chr8:<br>118 Mb | *Ngp* | chr8:118664419 | c.316C>T | p.Gln106* | stop_gained | HIGH | TT | CC | CC | CC | CC | CC | CC | CC | none |

**Figure 3.**
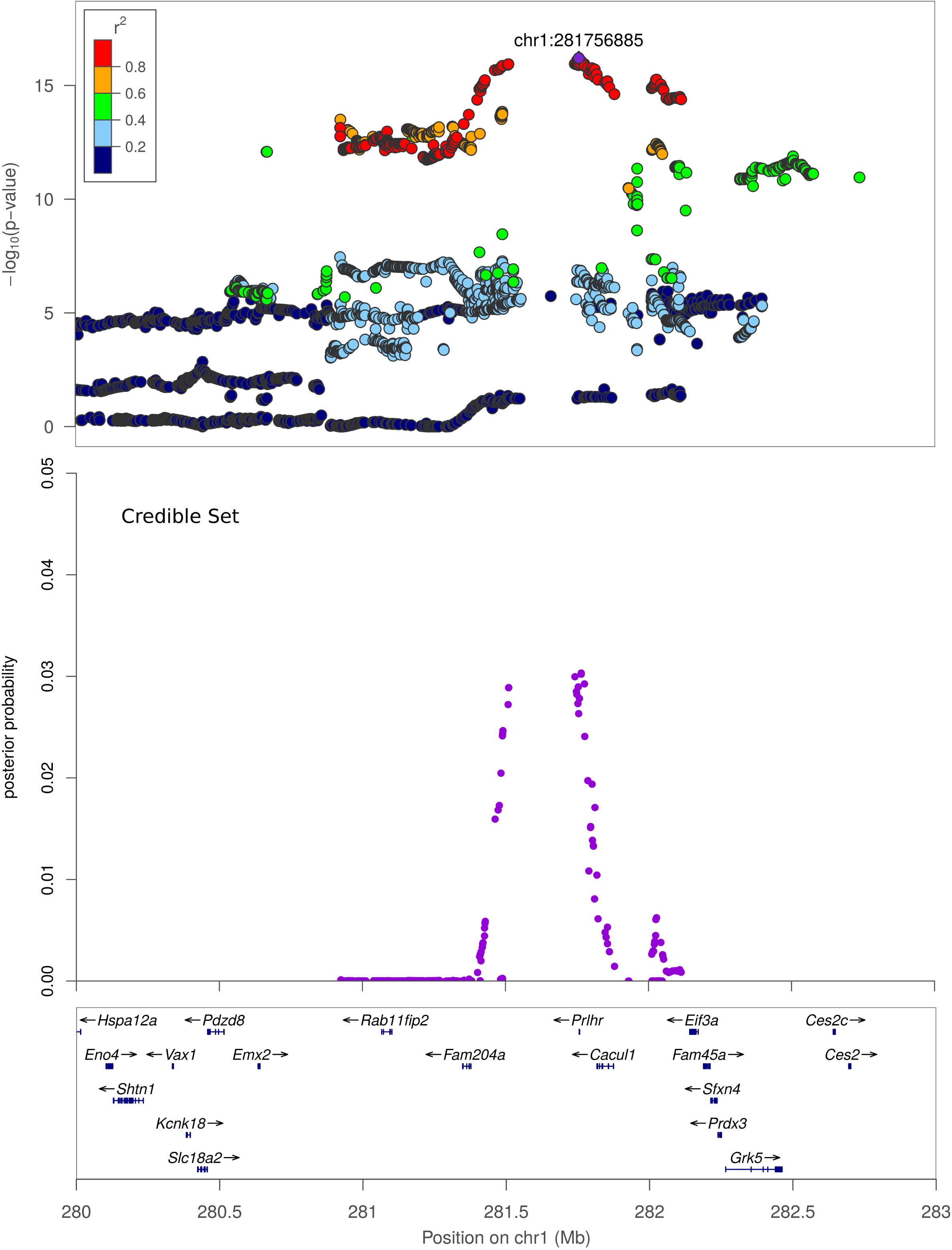
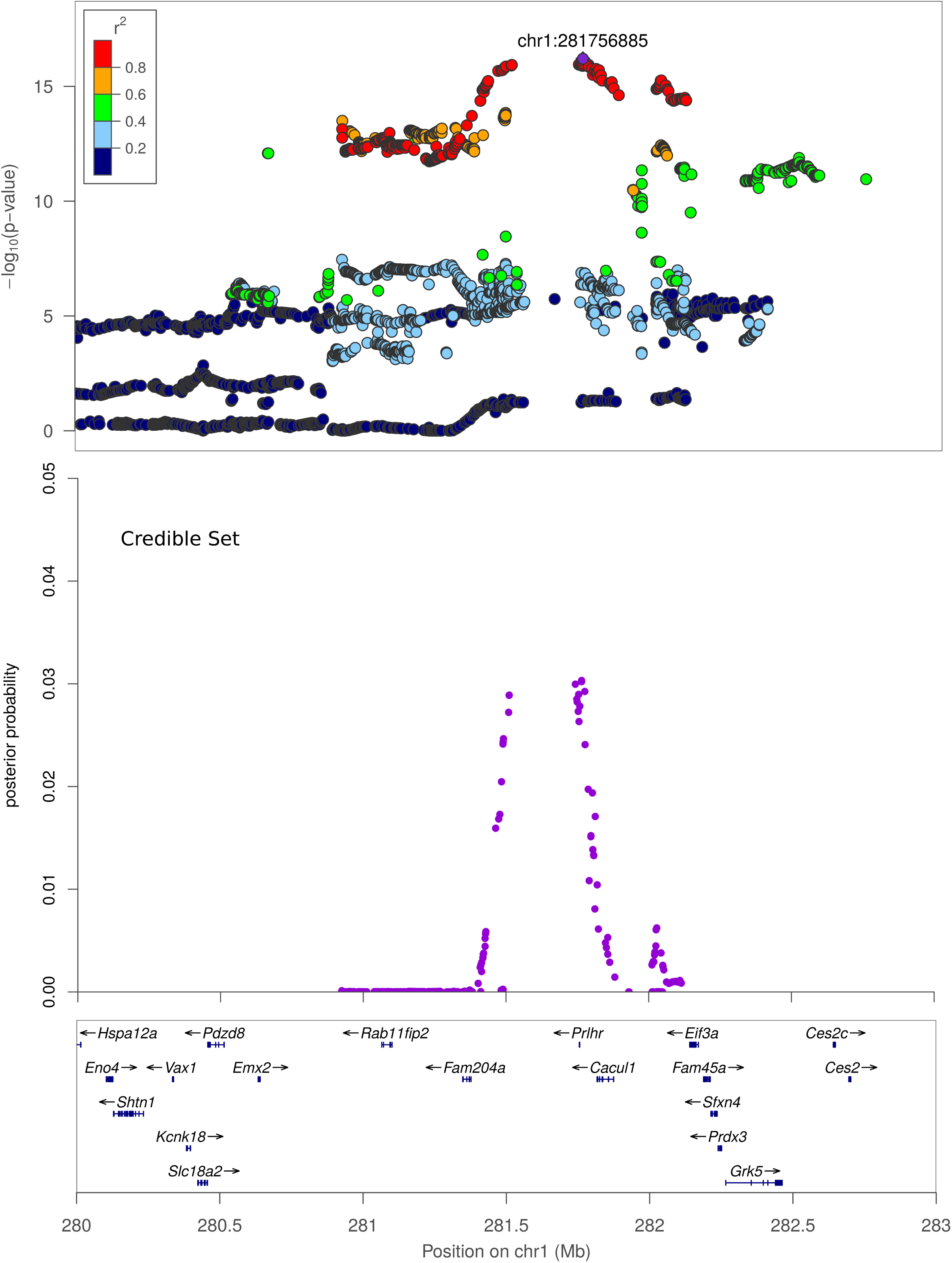
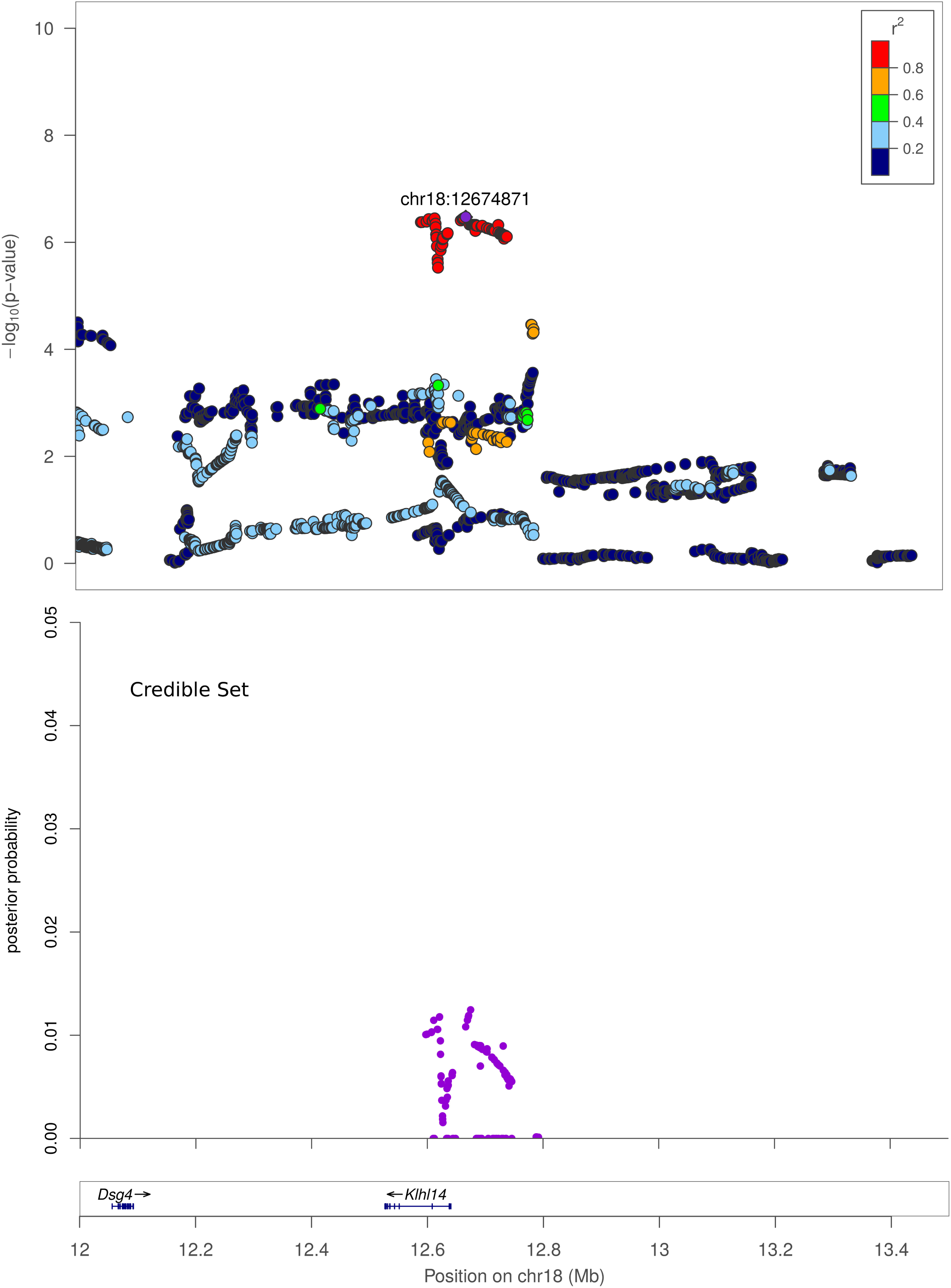
Regional association plots. X-axis shows chromosomal position. Top panel: Regional association plot. The SNPs with the lowest p-value (“top SNP”) is shown in purple, and its chromosomal position is indicated on the plot. Color of the dots indicates the level of linkage disequilibrium (LD) of each SNP with the top SNP. Middle panel: Credible set track shows posterior probability for being causal for the SNPs which were identified as the smallest set of SNPs accounting for 99% of the posterior probability (see Methods). Bottom panel: Genes in the region, as annotated by the Refseq. **A**. Regional association plot for Body weight at Chromosome 1, 280 Mb to 283 Mb. **B**. Regional association plot for Body Length_NoTail at Chromosome 16, 62.5 Mb to 65 Mb. **C**. Regional association plot for Body Length_NoTail at Chromosome 18, 12 Mb to 13.5 Mb.

## Discussion

The current study is the largest rodent GWAS ever reported. This study measured multiple adiposity traits collected at three different sites at multiple ages. Traits measured include body weight, body length, BMI (calculated with and without tail), fat pad weights and fasting glucose, all major risk factors for development of type 2 diabetes. We identified 32 significant loci, several of which correspond to narrow regions that contained logical candidate genes. We replicated previously identified loci and identified many novel loci. The large number of significant associations, the small regions implicated and the replication of previously reported loci despite age, diet and other environmental differences all highlight the power of HS rats for GWAS.

This study was part of a large multi-center behavioral study focused on genetic influences on drug abuse-relevant behaviors. As a result, the animals in this study were exposed to a variety of factors that may have altered adiposity. These include brief drug exposure, different behavioral experiences, as well as different diets and ages when tissues were collected. Despite these differences, we were successful at identifying a large number of adiposity QTL, indicating that these loci are resistant to multiple environmental influences. Importantly, contribution of these environmental influences, were taken into account in the statistical analysis by quantile normalizing data from each institution individually prior to running GWAS on the combined data. Although this design may have diluted our power to detect loci that were only relevant in one set of conditions, the loci that we did detect are likely to be robust to subtle drug, age and environmental differences. We view this aspect of the study as a strength as it allows us to generalize our findings across populations, including to humans in which there are multiple uncontrolled environmental influences

With the exception of fasting glucose, all traits demonstrate strong phenotypic and genetic correlations indicating that they are caused by overlapping sets of alleles. The pleiotropic loci that we identified further underscore the common genetic basis of these traits. The strongest pleiotropic locus identified falls on chr 1: 282 Mb and maps five of the adiposity traits,, with a second locus for RetroFat and EpiFat at chr. 6: 28 Mb, both replicating previous work by our group (Keele et al. 2018) (8) We also identified novel pleiotropic loci for body weight and body length (with and without tail) on chromosomes 3 and 7. Interestingly, we found that not all of the loci that contained multiple traits should be considered pleiotropic. We identified two regions that map more than one trait, but because both the mean allele frequency and the founder SDP do not match between the traits, these loci are likely driven by different genes and/or variants.

One of the strengths of using a highly recombinant outbred strain such as the HS is the ability to map to relatively small regions of the genome, greatly narrowing the number of potential candidate genes that may drive the QTL. In the current work, three of the identified loci contained only a single gene. For example, *EPH receptor A5 (Epha5)* is the only gene within the chr 14 locus for body length_NoTail. Ephrin receptors make up the largest sub-family of the receptor protein tyrosine kinases and are known to be involved in embryonic development, cell migration and axon guidance. Although *Epha5* has not previously been associated with body weight or height in human GWAS, *Epha5* knock-out mice have increased body weight relative to wild-type mice (24), suggesting a potential role for this gene in determining body length. In addition, *Neureglin1 (Nrg1)* is the only gene that falls within the chr 16 locus for body length_NoTail. *Nrg1* mediates cell-cell signaling and plays a critical role in growth and development of multiple organ systems. ICV injection of *Nrg1* leads to increased food intake and weight gain in rodents (25) and a recent study has demonstrated positive metabolic effect of *Nrg1* on multiple metabolic parameters, including body weight (26). In addition, *Nrg1* has been associated with BMI in a Korean population (27) making *Nrg1* a highly plausible gene within this region. Finally, *Kelch like family member 14 (Klhl14)* is the only gene that falls within the Epifat locus at chr 18. This gene localizes to the endoplasmic reticulum and has not previously been associated with adiposity traits in human or rat GWAS, suggesting that this finding could offer a novel biological insight. Regions containing a single gene are attractive, but we note that these regions may also contain regulatory variants for neighboring genes or un-annotated genes or transcripts that cause the association.

Another strength of using the HS is the ability to use the founder sequence to identify potentially damaging coding variants within QTL. We identified 154 coding variants, 63 of which match the SDP of the founder strains, making them plausible candidates. Replicating previous findings (8) we identified a high impact variant within *Prlhr*, a gene that falls within the highly significant pleiotropic locus on chr. 1, and which has previously been shown to play a role in feeding behavior (28, 29). We also identified a variant within *Adcy3*, a gene that falls within the chr. 6 locus for RetroFat and Epifat, and has been identified in both the HS rat (8) and human GWAS (30–32). We previously demonstrated that multiple genes likely play a role at the chr. 6 locus including *Adcy3, Krtcap3* and *Slc30a3* (see Keele et al. (8)). In addition to identifying candidate genes within these pleiotropic loci, we identified candidates within the fasting glucose loci: coding variants that match the SDP were identified in *Rnf213* (chr. 10 locus), a gene shown to protect beta cells and improve glucose tolerance (33), and *Xbp1* (chr. 14 locus), a gene shown to have multiple roles in glucose regulation (34). Finally, within the Body lenth_Tail QTL on chr. 8 we identified a high impact variant in *Ngp*, a gene that has not previously been associated with height or body weight. Future work will investigate the role of these variants in modifying adiposity traits.

This is the largest GWAS ever performed in rodent models and has identified a large number of significant loci. In spite of our large sample size, however, this study has its limitations. Adiposity measures were collected from both males and females, however, we have not presented sex-specific GWAS because the heritabilities for the two sexes were similar and because the reduction in sample size dramatically reduced the number of genome wide significant results. Therefore, we quantile normalized the males and females separately and then pooled them, which effectively removed mean differences between the sexes but did not account for gene by sex interactions. We are continuing to collect adiposity traits in additional animals and plan to use an even larger sample sizes to test for sex-specific QTLs in the future. A second limitation is that we have not incorporated expression QTL (eQTL) data for some of the most relevant tissues (e.g. liver and fat pads), which would be helpful for identifying candidate genes within the physiological QTL based on gene expression differences. Our study did not employee haplotype-based analyses, which is often used in highly recombinant populations such as the HS. Our laboratory is currently addressing several issues which may impact haplotype-based analysis including generating the genetic relationship matrix and the null distribution, as well as the most appropriate tool for determining founder probabilities. Finally, our study was part of a large multi-center behavioral study focused on genetic influences on drug abuse-relevant behaviors. As a result, a subset of the animals were briefly exposed to moderate doses of cocaine or nicotine, both of which could have changed body weight. In addition, the diets, ages and behavioral experiences of the rats differed across the three study sites. Any contribution of these environmental influences, however, were taken into account by including institution as a co-variate in the analysis. Although this design may have diluted our power to detect loci that were only relevant in one set of conditions, the loci that we did detect are likely to be robust to subtle drug, age and environmental differences, which can be viewed as a strength if the goal is to generalize our findings across populations, including to humans (35).

## Conclusions

The current study is the largest GWAS using a rodent population ever performed. We replicated previously identified loci from a smaller GWAS and identified numerous novel loci for multiple adiposity traits. Three of these loci contain only a single gene. Several other loci contain only a few genes, which simplifies the identification of candidate genes. We used prediction tools and founder sequence to identify candidate variants and genes within five of these loci. This work demonstrates the power of HS rats for fine-mapping and gene identification of adiposity traits, including the power to identify genetic loci across multiple institutions and environmental influence. This work also provides immediate candidate genes for future functional studies.

## Disclosure

The authors declared no conflict of interest.

## Notes

Funding: This work was supported by the National Institute on Drug Abuse (P50 DA037844) and the National Institute of Diabetes and Digestive and Kidney Diseases (R01 DK106386)

### Competing Interest Statement

The authors have declared no competing interest.

### Summary of Updates

Figure 2 revised; Figure 3 revised;

