## Supplementary material for "Genome wide association study in 3,173 outbred rats identifies multiple loci for body weight, adiposity, and fasting glucose": Supplemenatry material

**Supplementary Table S1**

**Behavioral testing at each phenotyping site**

Behavioral tests at NY site

| Approximate age | Procedure |
|---|---|
| 21 | Rats arrive in NY (approximately 21 days old). |
| 35 | Rats housed until they are approximately 35 days of age. |
| 49 | Social behavior |
| 51 | Locomotor activity. |
| 71 | Light reinforcement |
| 98 | Reaction Time |
| 120 | Delay Discounting: |
| 122 | Rats are transferred to a different animal facility |
| 127 | Pavlovian Condition Approach |
| 138 | Conditioned cue preference |
| 148 | Rats sacrificed; DNA for genotyping and tissue samples collected. |

Behavioral tests at MI site

| Approximate age | Procedure |
|---|---|
| 21 | Rats arrive in MI (approximately 21 days old). |
| 35 | Rats housed until they are approximately 35 days of age. |
| 42 | Pavlovian Condition Approach |
| 55 | Novelty seeking |
| 60 | Cocaine contextual conditioning |
| 75 | Rats sacrificed; DNA for genotyping and tissue samples collected. |

Behavioral tests at TN site

Breeders

| Approximate age | Procedure |
| --- | --- |
| 21 | Rats arrive in TN (approximately 21 days old). |
| 35 | Rats housed until breeding age and then bred for 1-2 cycles |
| 170 | Rats sacrificed; DNA for genotyping and tissue samples collected. |

Experimental rats

| Approximate age | Procedure |
| --- | --- |
| 0 | Rats are born from the HS breeders in TN animal facility |
| 21 | Behavior test battery (locomotor, social, etc) |
| 36 | Intravenous catheter implanter |
| 38 | Nicotine self-administration test for 12 days |
| 51 | Progressive ratio |
| 52 | Extinction |
| 55 | Reinstatement |
| 85 | Rats sacrificed; DNA for genotyping and tissue samples collected. |

**Supplementary Table S2**

Effect of a covariate was regressed out if it was significant and explained more than 2% of the variance

| Phenotype | covariates |
|---|---|
| Body Weight | Age, MI<br>Age, TN breeders<br>Age, TN experiment |
| Body Length NoTail | Age, NY<br>Age, TN breeders<br>Age, TN experiment<br>Technician JS, TN experiment<br>Technician XH, TN breeders |
| Body Length Tail | Age, MI<br>Age, NY<br>Age, TN experiment<br>Technician ALE, MI<br>Technician MM, MI<br>Technician YL, TN experiment |
| BMI NoTail | Age, MI<br>Age, NY<br>Age, TN experiment<br>Technician JS, TN experiment<br>Technician XH, TN breeders |
| BMI Tail | Age, MI<br>Age, NY<br>Age, TN breeders<br>Age, TN experiment<br>Technician ALE, MI<br>Technician MM, MI |
| RetroFat | Age, TN breeders<br>Age, TN experiment<br>Technician ALE, MI<br>Technician MM, MI |
| EpiFat | Age, TN breeders<br>Age, TN experiment<br>Technician AG, NY |

| | |
|---|---|
| | Technician CTA, NY<br>Technician NPR, NY |
| ParaFat | Age, NY<br>Age, TN experiment<br>Technician AH, MI<br>Technician KP, NY<br>Technician NPR, NY<br>Technician WH, TN breeders<br>Technician TW, TN experiment<br>Technician WH, TN experiment |
| Fasting Glucose | Age, NY |

**Supplementary Table S3**

**Sex specific SNP Heritability estimates**

| Trait | Males Heritability ± SE | Females Heritability ± SE |
|---|---|---|
| body weight | 0.45 ± 0.04 | 0.47 ± 0.04 |
| body length_Tail | 0.36 ± 0.04 | 0.35 ± 0.04 |
| body length_NoTail | 0.26 ± 0.04 | 0.27 ± 0.04 |
| BMI_Tail | 0.34 ± 0.04 | 0.25 ± 0.04 |
| BMI_NoTail | 0.28 ± 0.04 | 0.18 ± 0.04 |
| RetroFat | 0.47 ± 0.04 | 0.43 ± 0.04 |
| EpiFat | 0.37 ± 0.03 | NA |
| ParaFat | NA | 0.38 ± 0.04 |
| Fasting Glucose | 0.21 ± 0.04 | 0.19 ± 0.04 |

**Supplementary Table S4**

Summary of QTLs with strain distribution pattern of the founder strains. Pleiotropic loci are highlighted in yellow. Nearby loci that were not considered pleotropic are highlighted in grey.

| Trait | Chr | Peak Marker (bp) | -logP | Peak Marker | | | Genotype of the founders at the Peak Marker | | | | | | | | LD Interval | | | Genes in the LD interval (no LOC... ) | Credible Set Interval | | |
|---|---|---|---|---|---|---|---|---|---|---|---|---|---|---|---|---|---|---|---|---|---|
| | | | | Reference Allele | Allele frequency | Effect size | ACI | BN | BUF | F344 | M520 | MR | WN | WKY | start (bp ) | stop (bp ) | size (Mb) | | start (bp ) | stop (bp ) | size (Mb) |
| BMI_Tail | 1 | 106,866,154 | 6.05 | G | 0.29 | 0.15 ± 0.03 | A A | G G | G G | G G | G G | G G | G G | A A | 105,730,059 | 109,396,142 | 3.67 | *Ano5, Gas2, Ccdc179, Nell1, Fancf, Slc17a6, Svip* | 105,730,059 | 108,187,807 | 2.46 |
| RetroFat | 1 | 160,530,456 | 7.51 | T | 0.53 | -0.14 + 0.02 | C C | T T | T T | T T | T T | T T | C C | T T | 157,254,290 | 162,857,247 | 5.60 | *Dlg2, Ccdc90b, Mir708, Tenm4, Nars2, Gab2, Kctd21, Alg8, Ndufc2, Thrsp, Ints4, Aamdc, Clns1a, Aqp11, Pak1* | 157,254,290 | 162,857,247 | 5.60 |
| Body Weight | 1 | 185,730,317 | 7.58 | C | 0.74 | 0.16 ± 0.02 | T T | C C | T T | T T | C C | C C | C C | C C | 184,463,432 | 187,738,111 | 3.27 | *Xylt1, Sox6, Pik3c2a, Rps13, Plekha7, Nucb2* | 184,812,645 | 187,300,775 | 2.49 |
| BMI_NoTail | 1 | 187,300,775 | 6.72 | A | 0.66 | 0.15 ± 0.02 | G G | A A | G G | G G | G G | A A | A A | G G | 184,772,656 | 189,346,447 | 4.57 | *Gp2, Xylt1, Rps15a, Gde1, Itpripl2, Sox6, Umod, Acsm2, Tmc7, Iqck, Acsm5, Pik3c2a, Coq7, Syt17, Gprc5b, Gpr139, Knop1, Pdilt, Ccp110, Rps13, Clec19a, Smg1, Plekha7, Tmc5, Nucb2, Arl6ip1, Vps35l* | 184,816,720 | 189,346,447 | 4.53 |
| TL | 1 | 253,524,003 | 5.76 | G | 0.45 | 0.13 ± 0.03 | C C | G G | G G | G G | G G | G G | G G | C C | 253,082,171 | 254,725,734 | 1.64 | *Ears2l1,Rpp30,Kif20b,Pank1,Mir107,Htr7* | 253,376,067 | 254,725,734 | 1.35 |
| ParaFat | 1 | 280,924,549 | 7.22 | G | 0.57 | 0.20 ± 0.03 | G G | G G | A A | G G | G G | G G | G G | A A | 280,876,316 | 282,114,080 | 1.24 | *Fam204a, Cacul1, Grk5, Prlhr, Eif3a, Ces2i, Nanos1, Prdx3, Rab11fip2* | 280,924,333 | 282,114,080 | 1.19 |
| Body Weight | 1 | 281,756,885 | 16.21 | C | 0.58 | 0.21 ± 0.02 | C C | C C | T T | C C | C C | C C | C C | T T | 280,924,333 | 282,114,080 | 1.19 | *Fam204a, Prlhr, Cacul1* | 281,402,451 | 282,114,080 | 0.71 |
| RetroFat | 1 | 281,777,218 | 20.21 | A | 0.57 | 0.24 + 0.02 | A A | A A | C C | A A | A A | A A | A A | C C | 280,924,333 | 282,114,080 | 1.19 | *Fam204a, Cacul1, Grk5, Prlhr, Eif3a, Ces2i, Nanos1, Prdx3, Rab11fip2* | 280,924,333 | 282,043,175 | 1.12 |
| EpiFat | 1 | 281,802,657 | 14.00 | C | 0.56 | 0.29 ± 0.03 | C C | C C | T T | C C | C C | C C | **T T** | T T | 280,924,333 | 282,736,277 | 1.81 | *Fam45a, Fam204a, Cacul1, Grk5, Prlhr, Eif3a, Ces2i, Nanos1, Prdx3, Rab11fip2, Ces2, Ces2c, Sfxn4* | 280,924,333 | 282,114,080 | 1.19 |
| BMI_Tail | 1 | 282,049,439 | 11.44 | C | 0.55 | 0.18 ± 0.02 | C C | C C | T T | C C | C C | C C | C C | T T | 280,924,333 | 282,736,277 | 1.81 | *Fam45a, Fam204a, Cacul1, Grk5, Prlhr, Eif3a, Ces2i, Nanos1, Rab11fip2, Prdx3, Ces2, Ces2c, Sfxn4* | 280,924,333 | 282,114,080 | 1.19 |
| Body Weight | 2 | 65,816,485 | 6.55 | A | 0.91 | 0.20 ± 0.03 | G G | A A | G G | G G | G G | G G | G G | G G | 62,570,942 | 71,814,490 | 9.24 | *Cdh10, Cdh9, Cdh12, Drosha, Cdh6, Egf* | 62,570,942 | 71,814,490 | 9.24 |
| RetroFat | 3 | 95,389,621 | 13.12 | A | 0.44 | -0.19 + 0.02 | A A | A A | A A | A A | G G | A A | G G | A A | 92,336,188 | 97,685,154 | 5.35 | Cd59, Ccdc73, Prrg4, Apip, Qser1, Cstf3, Nat10, Pdhx, Immp1l, Tcp11l1, Eif3m, Cat, Pamr1, Dcdc1, Abtb2, Pax6, Rcn1, Ehf, Caprin1, Dcdc5, Lmo2, Depdc7, Elf5, Dnajc24, Hipk3, Wt1, Slc1a2, Cd44, Fbxo3, Elp4 | 94,050,143 | 95,685,634 | 1.64 |
| Body Length_NoTail | 3 | 136,021,511 | 6.92 | A | 0.77 | -0.16 ± 0.03 | G G | A A | G G | G G | G G | G G | G G | G G | 132,291,573 | 137,146,532 | 4.85 | *Sel1l2, Ism1, Flrt3, Snrpb2, Tasp1, Esf1, Macrod2, Sptlc3, Ndufaf5, Kif16b* | 132,865,862 | 136,625,612 | 3.76 |
| Body Length_Tail | 3 | 136,021,511 | 5.97 | A | 0.77 | -0.14 ± 0.03 | G G | A A | G G | G G | G G | G G | G G | G G | 132,291,573 | 137,146,532 | 4.85 | *Sel1l2, Ism1, Flrt3, Snrpb2, Tasp1, Esf1, Macrod2, Ndufaf5, Sptlc3, Kif16b* | 132,779,438 | 137,146,532 | 4.37 |
| Body Weight | 3 | 136,021,511 | 7.34 | A | 0.77 | -0.16 ± 0.02 | G G | A A | G G | G G | G G | G G | G G | G G | 132,291,573 | 137,146,532 | 4.85 | *Sel1l2, Ism1, Flrt3, Tasp1, Esf1, Macrod2, Snrpb2, Ndufaf5, Sptlc3, Kif16b* | 134,319,381 | 137,146,532 | 2.83 |
| RetroFat | 3 | 137,537,161 | 8.21 | G | 0.44 | -0.15 + 0.02 | A A | G G | G G | G G | A A | G G | G G | G G | 136,161,761 | 138,849,437 | 2.69 | Bfsp1, Banf2, Dstn, Rbbp9, Kat14, Otor, Polr3f, Smim26, Macrod2, Snrpb2, Dzank1, Mgme1, Pcsk2, Zfp133, Dtd1, Ovol2, Kif16b, Rrbp1, Snx5, Sec23b | 136,178,205 | 138,775,911 | 2.60 |
| Body Weight | 5 | 50,933,779 | 6.99 | A | 0.78 | -0.16 ± 0.03 | G G | G G | G G | G G | A A | G G | G G | G G | 49,152,709 | 50,940,275 | 1.79 | *Smim8, Ifnk, Cga, Spaca1, Orc3, Akirin2, Slc35a1, Cfap206, Cnr1, Rars2, Mob3b, Zfp292* | 49,156,473 | 50,940,275 | 1.78 |
| EpiFat | 6 | 26,266,960 | 7.25 | G | 0.59 | -0.20 ± 0.03 | A A | G G | G G | G G | G G | G G | G G | A A | 22,684,886 | 28,223,800 | 5.54 | *Alk, Clip4, Wdr43, Spdya, Ppp1cb, Srd5a2, Plb1, Fosl2, Babam2, Rbks, Slc4a1ap, Supt7l, Gpn1, Gckr, Ift172, Krtcap3, Nrbp1, Ppm1g, Zfp513, Snx17, Eif2b4, Gtf3c2, Mpv17, Ucn, Trim54, Dnajc5g, Slc30a3, Cad, Atraid, Slc5a6, Prr30, Preb, Abhd1, Cgref1, Khk, Emilin1, Ost4, Agbl5, Tmem214, Mapre3, Dpysl5, Cenpa, Slc35f6, Kcnk3, Otof, Drc1, Selenoi, Adgrf3, Hadhb, Hadha, Garem2, Rab10, Kif3c, Dtnb* | 22,940,602 | 28,164,637 | 5.22 |
| RetroFat | 6 | 28,148,338 | 19.50 | G | 0.65 | -0.26 + 0.02 | **G G** | G G | G G | G G | G G | G G | G G | T T | 25,954,450 | 28,752,109 | 2.80 | Cenpo, Asxl2, Zfp512, Krtcap3, Agbl5, Tcf23, Dnajc27, Zfp513, Tmem214, Trnaa-agc6, Emilin1, Slc35f6, Mapre3, Drc1, Preb, Rab10, Ppm1g, Trim54, Snx17, Gpn1, Nrbp1, Ncoa1, Prr30, Trnay-gua3, Adcy3, Atraid, Abhd1, Khk, Slc5a6, Cib4, Ost4, Cad, Cenpa, Kif3c6 Rbks, Efr3b, Gckr, Gtf3c2, Dnmt3a, Ucn, Eif2b4, Hadhb, Pomc, Ptrhd1, Slc4a1ap, Dtnb, Adgrf3, Slc30a3, Selenoi, Mrpl33, Otof, Babam2, Hadha, Ift172, Dnajc5g, Kcnk3, Garem2, Supt7l | 26,844,333 | 28,707,142 | 1.86 |
| Body Length_Tail | 6 | 137,745,191 | 5.89 | C | 0.54 | -0.12 + 0.02 | T T | C C | C C | C C | C C | C C | C C | T T | 136,769,305 | 138,087,183 | 1.32 | *Tmem179, Mta1, Cdca4, Crip2, Ahnak2, Brf1, Cep170b, Adssl1, Gpr132, Clba1, Pld4, Tedc1, Siva1, Tmem121, Akt1, Zbtb42, Ighe, Nudt14, Tex22, Inf2, Jag2, Pacs2, Btbd6, Crip1* | 137,264,981 | 138,087,183 | 0.82 |
| Body Weight | 7 | 24,886,476 | 7.29 | G | 0.07 | 0.26 ± 0.04 | G G | G G | C C | G G | G G | G G | G G | G G | 24,869,890 | 25,213,445 | 0.34 | *Tcp11l2, Nuak1, Ckap4* | 24,869,890 | 25,144,762 | 0.27 |
| Body Weight | 7 | 36,497,588 | 9.77 | C | 0.14 | -0.24 ± 0.03 | C C | C C | **G G** | C C | G G | G G | C C | C C | 34,119,928 | 36,522,260 | 2.40 | *Usp44, Rpl13l3, Hal, Mir331, Vezt, Ndufa12, Plxnc1, Snrpf, Elk3, Lta4h, Ntn4, Trnad-guc2, Cradd, Amdhd1, Fgd6, Tmcc3, Socs2, Metap2, Nr2c1, Ccdc38, Cep83* | 34,156,704 | 36,517,726 | 2.36 |
| Body Length_NoTail | 7 | 36,517,726 | 7.88 | T | 0.12 | -0.25 ± 0.04 | T T | T T | T T | T T | C C | C C | T T | T T | 34,119,928 | 36,522,260 | 2.40 | *Lta4h, Hal, Amdhd1, Snrpf, Ntn4, Usp44, Metap2, Mir331, Vezt, Fgd6, Nr2c1, Ndufa12, Tmcc3, Cradd, Socs2* | 34,156,704 | 36,522,260 | 2.37 |
| Body Length_Tail | 7 | 36,517,726 | 14.37 | T | 0.12 | -0.33 ± 0.04 | T T | T T | T T | T T | C C | C C | T T | T T | 34,119,928 | 36,522,260 | 2.40 | *Hal, Rpl13l3, Usp44, Mir331, Vezt, Snrpf, Plxnc1, Ndufa12, Elk3, Lta4h, Ntn4, Trnad-guc2, Cradd, Fgd6, Amdhd1, Metap2, Socs2, Nr2c1, Tmcc3, Ccdc38, Cep83* | 34,903,746 | 36,522,260 | 1.62 |
| TL | 7 | 36,526,715 | 6.10 | A | 0.37 | 0.14 ± 0.03 | A A | A A | A A | A A | A A | A A | A A | G G | 36,289,478 | 36,765,903 | 0.48 | *Nudt4,Mrpl42,Ube2n,Socs2,Cradd* | 36,349,309 | 36,554,099 | 0.20 |
| Body Length_Tail | 8 | 118,711,320 | 5.72 | A | 0.67 | 0.13 ± 0.02 | C C | A A | A A | A A | A A | A A | A A | A A | 116,614,891 | 119,781,444 | 3.17 | *Pth1r, Myl3, Actl11, Usp19, Wdr6, Camp, Setd2, Prss46, Prss45, Qrich1, Gpx1, Cdc25a, Mst1, Pfkfb4, Nme6, Celsr3, Ptpn23, Inka1, Qars, Klhdc8b, P4htm, Epm2aip1, Slc26a6, Map4, Ltf, Lrrfip2, Traip, Mir3555, Mir191, Trank1, Ccdc71, Prss44, Rbm6, Mst1r, Fbxw12, Ccdc12, Mon1a, Prss50, Ip6k1, Rtp3, Ndufaf3, Cspg5, Impdh2, Smarcc1, Ccdc51, Trex1, Ccrl2, Atrip, Tdgf1, Gmppb, Prss42, Arih2os, Apeh, Uba7, Als2cl, Klhl18, Amigo3, Dhx30, Ngp, Dclk3, Nicn1, Kif9, Rnf123, Mir425, Mlh1, Ip6k2, Rhoa, Camkv, Bsn, Nradd, Arih2, Nbeal2, Cdhr4, Elp6, Ccdc36, Prkar2a, Dalrd3, Amt, Nckipsd, Usp4, Dag1, Scap, Lrrc2, Spink8, Uqcrc1, Mir711, Ucn2, Col7a1, Shisa5, Tmie, Lamb2, Slc25a20, Tma7, Plxnb1, Tcta, Tmem89* | 117,632,634 | 119,768,313 | 2.14 |
| Body Length_Tail | 9 | 65,078,205 | 6.81 | C | 0.76 | 0.16 ± 0.03 | T T | C C | T T | C C | T T | T T | T T | T T | 64,013,585 | 65,670,399 | 1.66 | *Ndufb3, Aox4, Aox3, Casp8, Als2cr12, Tyw5, Mars2, Orc2, Kctd18, Aox1, Ppil3, Fam126b, Aox2, Sgo2, Clk1, Maip1, Spats2l, Cflar, Nif3l1, Bzw1* | 64,063,377 | 65,670,399 | 1.61 |
| BMI_Tail | 10 | 84,080,794 | 8.21 | G | 0.57 | -0.16 ± 0.02 | A A | G G | A A | G G | A A | G G | G G | G G | 83,442,285 | 85,006,252 | 1.56 | *Snf8, Hoxb4, Osbpl7, Hoxb9, Mir10a, Cdk5rap3, Hoxb5os, Atp5mc1, Scrn2, Phb, Sp2, Mir196c, Hoxb5, Hoxb7, Hoxb6, Gngt2, Pnpo, Hoxb13, Gip, B4galnt2, Ttll6, Ube2z, Lrrc46, Nfe2l1, Abi3, Prr15l, Hoxb2, Hoxb8, Mrpl10, Hoxb3, Skap1, Hoxb1, Snx11, Mir152, Zfp652, Phospho1, Copz2, Trnaq-uug1, Igf2bp1, Cbx1, Sp6, Calcoco2* | 83,442,285 | 84,604,288 | 1.16 |
| TL | 10 | 84,263,936 | 9.38 | T | 0.57 | 0.18 ± 0.03 | C C | T T | C C | T T | C C | T T | T T | T T | 83,442,285 | 85,006,252 | 1.56 | *Snf8, Hoxb4, Osbpl7, Hoxb9, Mir10a, Cdk5rap3, Hoxb5os, Atp5mc1, Scrn2, Phb, Sp2, Mir196c, Hoxb5, Hoxb7, Hoxb6, Gngt2, Pnpo, Hoxb13, Gip, B4galnt2, Ttll6, Ube2z, Lrrc46, Nfe2l1, Abi3, Prr15l, Hoxb2, Hoxb8, Mrpl10, Hoxb3, Skap1, Hoxb1, Snx11, Mir152, Zfp652, Phospho1, Copz2, Trnaq-uug1, Igf2bp1, Cbx1, Sp6, Calcoco2* | 83,777,266 | 84,604,288 | 0.83 |
| Body Length_Tail | 10 | 85,082,795 | 7.37 | C | 0.72 | 0.17 ± 0.03 | G G | C C | G G | C C | G G | C C | C C | G G | 84,902,901 | 85,239,899 | 0.34 | *Osbpl7, Tbx21, Npepps, Sp2, Scrn2, Lrrc46, Mrpl10, Tbkbp1, Kpnb1, Sp6* | 85,063,786 | 85,239,899 | 0.18 |
| BMI_NoTail | 10 | 96,804,258 | 7.66 | T | 0.48 | -0.16 ± 0.02 | C C | T T | C C | T T | C C | T T | T T | C C | 96,561,667 | 98,097,621 | 1.54 | *Prkca, Apoh, Cep112, Axin2, Rpl-1, Mif, Rgs9, Gna13, Gpa-12, Amz2, Slc16a6, Arsg, Wipi1, Prkar1a, Kin-2, Fam20a* | 96,561,667 | 98,097,621 | 1.54 |
| Fasting Glucose | 10 | 109,944,213 | 6.33 | A | 0.75 | -0.18 ± 0.03 | T T | A A | T T | T T | T T | T T | T T | T T | 108,350,175 | 110,315,359 | 1.97 | *Ccdc40, Gaa, Eif4a3, Nptx1, Rptor, Chmp6, Baiap2, Aatk, Mir3065, Mir338, Cep131, Mir3594, Mir3562, Tepsin, Ndufaf8, Actg1, Fscn2, Nploc4, Oxld1, Ccdc137, Hgs, Mrpl12, Slc25a10, Gcgr, Mcrip1, Ppp1r27, P4hb, Arhgdia, Alyref, Anapc11, Npb, Pcyt2, Sirt7, Mafg, Pycr1, Myadml2, Notum, Lrrc45, Dcxr, Rfng, Gps1, Dus1l, Fasn, Slc16a3, Csnk1d, Cd7, Sectm1b, Sectm1a, Tex19.2* | 108,362,930 | 110,315,359 | 1.95 |
| Body Weight | 10 | 111,010,289 | 6.10 | T | 0.41 | 0.13 ± 0.02 | C C | T T | T T | C C | C C | C C | C C | C C | 110,664,101 | 111,022,246 | 0.36 | *Tbcd, Znf750, B3gntl1, Metrnl* | 110,912,475 | 111,022,246 | 0.11 |
| Body Length_Tail | 12 | 2,199,384 | 8.94 | C | 0.18 | -0.22 ± 0.03 | T T | C C | T T | C C | C C | C C | C C | T T | 846,435 | 5,587,624 | 4.74 | *Stxbp2, Kl, Snapc2, Rfc3, Higd2al1, Cd209f, Xab2, Cers4, Vom2r-ps90, Retn, Ctxn1, Dppa1-ps1, Camsap3, Elavl1, Pcp2, Lrrc8e, Evi5l, Cd209e, Vom2r60, Insr, Clec4g, Fcer2, Arhgef18, Prr36, Fry, Clec4m, Tex45, Rn5s, Trappc5, Vom2r-ps97, Vom2r58, Ccl25, Cd209c, Vom2r-ps91, Map2k7, Pet100, Cd209, Tgfbr3l, Cd209a, Mcoln1, Pnpla6, Timm44, Zfp358, Stard13, Vom2r-ps96, Vom2r-ps95, Vom2r-ps94, Mcemp1, Vom2r59, Pex11g* | 874,769 | 1,365,748 | 0.49 |
| Body Weight | 12 | 5,738,696 | 9.12 | C | 0.59 | 0.16 ± 0.02 | C C | C C | C C | T T | T T | T T | T T | C C | 4,938,015 | 6,078,451 | 1.14 | *Rxfp2, Zfp958, Fry, Vom2r58, Vom2r-ps91, Lnc001* | 5,557,587 | 6,054,407 | 0.50 |
| RetroFat | 12 | 5,782,829 | 7.12 | T | 0.73 | 0.14 + 0.02 | C C | T T | C C | C C | C C | C C | C C | C C | 455,837 | 6,259,634 | 5.80 | Rxfp2, Zar1l, Pds5b, N4bp2l2, Zfp958, Stxbp2, Kl, Snapc2, Rfc3, Higd2al1, Cd209f, Brca2, Xab2, Cers4, Vom2r-ps90, N4bp2l1, Retn, Ctxn1, Dppa1-ps1, Camsap3, Elavl1, Pcp2, Lrrc8e, Evi5l, Cd209e, Vom2r60, Insr, Clec4g, Fcer2, Arhgef18, Prr36, Fry, Clec4m, LOC688432, Tex45, Rn5s, Trappc5, Vom2r-ps97, Vom2r58, Ccl25, Cd209c, Vom2r-ps91, Map2k7, Pet100, Cd209, Tgfbr3l, Cd209a, Mcoln1, Pnpla6, Timm44, Zfp358, Stard13, Vom2r-ps96, Vom2r-ps95, Vom2r-ps94, Mcemp1, Vom2r59, Pex11g, Lnc001 | 455,837 | 6,259,634 | 5.80 |
| Body Length_NoTail | 12 | 6,239,515 | 6.26 | A | 0.52 | 0.13 ± 0.02 | A A | A A | A A | G G | G G | G G | G G | G G | 6,078,237 | 7,035,028 | 0.96 | *Uspl1, Rxfp2, Medag, Hsph1, Trnan-guu11, Tex26, B3glct, Alox5ap, Wdr95* | 6,078,237 | 7,035,028 | 0.96 |
| Body Length_Tail | 12 | 43,060,205 | 6.27 | G | 0.75 | -0.15 ± 0.02 | G G | G G | A A | A A | A A | A A | A A | A A | 42,024,283 | 43,190,584 | 1.17 | *Oas1g, Tbx3, Tbx5* | 42,062,846 | 43,172,537 | 1.11 |
| RetroFat | 13 | 55,021,887 | 7.16 | C | 0.67 | 0.14 + 0.02 | G G | C C | G G | C C | C C | G G | C C | C C | 54,910,667 | 56,487,438 | 1.58 | *Mir3570, Mir181, Ptprc, Atp6v1g3, Nek7, Lhx9, Crb1* | 54,910,667 | 56,387,208 | 1.48 |
| TL | 13 | 109,566,014 | 7.04 | T | 0.34 | 0.15 ± 0.03 | C C | T T | T T | T T | T T | C C | T T | T T | 108,055,241 | 111,868,706 | 3.81 | *Lpgat1,Slc30a1,Cenpf,Batf3,Hhat,Syt14,Ptpn14,Flvcr1,Tatdn3,Fam71a,Nenf,Nek2,Ppp2r5a,Rd3,Utp25,Prox1,Atf3,Rcor3,Sertad4,Rps6kc1,Angel2,Vash2,Nsl1,RGD1560186,Traf5,Smyd2,Pacc1,Dtl,Ints7,Kcnh1* | 108,055,241 | 110,873,866 | 2.82 |
| Body Length_NoTail | 14 | 26,365,986 | 6.54 | A | 0.73 | -0.16 ± 0.03 | G G | A A | G G | G G | G G | G G | G G | A A | 25,129,356 | 26,478,660 | 1.35 | *Epha5* | 25,725,806 | 26,478,660 | 0.75 |
| Fasting Glucose | 14 | 86,029,588 | 6.27 | T | 0.27 | -0.17 ± 0.03 | T T | T T | T T | T T | T T | T T | T T | C C | 82,473,632 | 87,575,807 | 5.10 | *Nefh, Dusp18, Znrf3, Rnf185, Patz1, Ccdc117, Kremen1, Tug1, Maea, Depdc5, Ddx56, Myl7, Pold2, Thoc5, Rhbdd3, Lif, Slc35e4, Sec14l4, Frg2, Ccdc157, Gck, Camk2b, Nudcd3, Pla2g3, Castor1, Aebp1, Adcy1, Mrps24, Cabp7, Tcn2, Urgcp, Purb, Nkx1-1, Limk2, Spon2, Rnf215, Pisd, Tbc1d10a, Slc5a1, Dbnl, Nipsnap1, Ewsr1, Myo1g, Uvssa, Hormad2, Smtn, Polm, Wap, Sec14l3, Gal3st1, Osm, Pik3ip1, Selenom, Snrpg-ps1, Uqcr10, Mtmr3, Zmat5, Ctbp1, Ppia, Igfbp1, Ywhah, Nf2, Ramp3, Sec14l2, H2afv, Ascc2, Tmed4, Morc2, Prr14l, Ykt6, Ap1b1, Nacad, Sfi1, Pes1, Ankrd36, Tbrg4, Ccm2, Zmiz2, Inpp5j, Xbp1, Emid1, Sf3a1, Npc1l1, Mtfp1, Pgam2, Igfbp3, Eif4enif1, Eif6-ps1, Ogdh, Drg1, Rasl10a, Gas2l1, Osbp2* | 85,206,177 | 86,459,891 | 1.25 |
| Body Length_NoTail | 16 | 64,014,119 | 6.35 | G | 0.55 | 0.13 ± 0.02 | G G | G G | G G | A A | G G | G G | A A | A A | 64,003,027 | 64,416,540 | 0.41 | *Nrg1* | 64,003,283 | 64,416,540 | 0.41 |
| EpiFat | 18 | 12,674,871 | 6.47 | G | 0.22 | -0.22 ± 0.04 | G G | G G | G G | G G | G G | G G | G G | A A | 12,597,378 | 12,792,162 | 0.19 | *Klhl14* | 12,597,378 | 12,745,926 | 0.15 |
| BMI_NoTail | 18 | 25,190,274 | 5.78 | T | 0.04 | -0.34 ± 0.07 | C C | T T | T T | T T | T T | C C | C C | T T | 23,785,532 | 25,418,376 | 1.63 | *Ercc3, Ammecr1l, Rit2, Potef, Tex51, Sft2d3, Ubbp4, Map3k2, Iws1, Lims2, Myo7b, Gpr17, Proc, Sap130, Slc25a46, Wdr33, Gypc, Polr2d, Bin1, Syt4* | 23,848,600 | 25,364,185 | 1.52 |
| TL | 19 | 32,004,685 | 7.81 | C | 0.41 | -0.15 ± 0.03 | C C | C C | C C | T T | T T | C C | C C | C C | 31,176,512 | 32,698,569 | 1.52 | *Anapc10,Mmaa,Zfp827,Abce1,Smad1,Otud4,Hhip* | 31,176,512 | 32,260,219 | 1.08 |

**Supplementary Table S5.** Variant annotation and SDP for 153 potentially damaging coding variants identified for 18 QTLs **.** Yellow highlights indicate coding variants in which the SDP matches that of the peak marker at the QTL.

| Trait | Peak.Marker | SNP in LD with Peak Marker | ref | alt | effect | impact | gene | Transcript.ID | cDNA.position | SNP.change | Amino.acid.change | ACI | BN | BUF | F344 | M520 | MR | WN | WKY |
|---|---|---|---|---|---|---|---|---|---|---|---|---|---|---|---|---|---|---|---|
| Fasting Glucose | chr10:109944213 | chr10:108582280 | C | G | missense_variant | MODERATE | *Rnf213* | ENSRNOT00000044983.5 | 9502 | c.9502C>G | p.His3168Asp | GG | CC | GG | GG | GG | GG | GG | GG |
| | | chr10:108582544 | T | C | missense_variant | MODERATE | *Rnf213* | ENSRNOT00000044983.5 | 9766 | c.9766T>C | p.Trp3256Arg | CC | TT | CC | CC | CC | CC | CC | CC |
| | | chr10:108604073 | G | C | missense_variant | MODERATE | *Rnf213* | ENSRNOT00000044983.5 | 12263 | c.12263G>C | p.Ser4088Thr | CC | GG | CC | CC | CC | CC | CC | CC |
| | | chr10:109244743 | T | C | missense_variant | MODERATE | *Cep131* | ENSRNOT00000005977.6 | 2802 | c.2729A>G | p.Asn910Ser | CC | TT | CC | CC | CC | CC | CC | CC |
| | | chr10:109251307 | A | G | missense_variant | MODERATE | *Cep131* | ENSRNOT00000005977.6 | 1311 | c.1238T>C | p.Leu413Pro | GG | AA | GG | GG | GG | GG | GG | GG |
| | | chr10:109264307 | T | C | missense_variant | MODERATE | *Cep131* | ENSRNOT00000005977.6 | 125 | c.52A>G | p.Met18Val | CC | TT | CC | CC | CC | CC | CC | CC |
| | | chr10:109286225 | G | A | missense_variant | MODERATE | *Slc38a10* | ENSRNOT00000006225.8 | 2491 | c.2231C>T | p.Ala744Val | AA | GG | AA | AA | AA | AA | AA | AA |
| | | chr10:109296990 | A | G | missense_variant | MODERATE | *Slc38a10* | ENSRNOT00000006225.8 | 1528 | c.1268T>C | p.Val423Ala | GG | AA | GG | GG | GG | GG | GG | GG |
| | | chr10:109549959 | C | G | missense_variant | MODERATE | *Faap100* | ENSRNOT00000054974.4 | 697 | c.695G>C | p.Ser232Thr | GG | CC | GG | GG | GG | GG | GG | GG |
| | | chr10:109550184 | T | C | missense_variant | MODERATE | *Faap100* | ENSRNOT00000054974.4 | 472 | c.470A>G | p.His157Arg | CC | TT | CC | CC | CC | CC | CC | CC |
| | | chr10:110252682 | T | C | missense_variant | MODERATE | *Sectm1b* | ENSRNOT00000054932.4 | 405 | c.235A>G | p.Lys79Glu | CC | TT | CC | CC | CC | CC | CC | CC |
| Body Length_Tail | chr12:2199384 | chr12:2001010 | C | A | missense_variant | MODERATE | *Pex11g* | ENSRNOT00000037564.4 | 569 | c.520G>T | p.Val174Leu | CC | CC | CC | AA | AA | AA | AA | CC |
| Tail Length | chr13:109566014 | chr13:108143993 | T | A | missense_variant | MODERATE | *Cenpf* | ENSRNOT00000004525.6 | 13 | c.6071A>T | p.Asn2024Ile | AA | TT | TT | TT | TT | AA | TT | AA |
| | | chr13:108147187 | C | T | missense_variant | MODERATE | *Cenpf* | ENSRNOT00000004525.6 | 12 | c.4382G>A | p.Gly1461Glu | TT | CC | CC | CC | CC | TT | CC | TT |
| | | chr13:108148751 | G | A | missense_variant | MODERATE | *Cenpf* | ENSRNOT00000004525.6 | 12 | c.2818C>T | p.Leu940Phe | AA | GG | GG | GG | GG | AA | GG | AA |
| | | chr13:108149351 | C | T | missense_variant | MODERATE | *Cenpf* | ENSRNOT00000004525.6 | 12 | c.2218G>A | p.Val740Ile | TT | CC | CC | CC | CC | TT | CC | TT |
| | | chr13:108444314 | A | G | missense_variant | MODERATE | *LOC498308* | ENSRNOT00000048200.1 | 7 | c.569T>C | p.Val190Ala | GG | AA | AA | AA | AA | GG | AA | AA |
| | | chr13:108444439 | T | G | missense_variant | MODERATE | *LOC498308* | ENSRNOT00000048200.1 | 6 | c.556A>C | p.Thr186Pro | GG | TT | TT | TT | TT | GG | TT | TT |
| | | chr13:108444440 | G | T | missense_variant | MODERATE | *LOC498308* | ENSRNOT00000048200.1 | 6 | c.555C>A | p.Asn185Lys | TT | GG | GG | GG | GG | TT | GG | GG |
| | | chr13:108681633 | A | G | missense_variant | MODERATE | *Smyd2* | ENSRNOT00000004783.6 | 9 | c.851A>G | p.Asn284Ser | GG | AA | AA | AA | AA | GG | GG | AA |
| | | chr13:110372281 | C | G | missense_variant | MODERATE | *RGD15601* | ENSRNOT00000049634.2 | 1 | c.118C>G | p.Pro40Ala | GG | CC | CC | CC | CC | GG | CC | CC |
| | | chr13:55284064 | C | T | missense_variant | MODERATE | *Atp6v1g3* | ENSRNOT00000029679.4 | 194 | c.166C>T | p.Arg56Cys | CC | CC | CC | TT | CC | CC | TT | CC |
| | | chr13:56262414 | C | T | missense_variant | MODERATE | *AABR0702* | ENSRNOT00000032908.2 | 209 | c.209C>T | p.Pro70Leu | TT | CC | TT | CC | TT | TT | CC | CC |
| RetroFat | chr13:55021887 | chr13:56262920 | G | A | missense_variant | MODERATE | *AABR0702* | ENSRNOT00000032908.2 | 692 | c.692G>A | p.Arg231His | GG | GG | GG | AA | GG | GG | AA | GG |
| | | chr13:56262923 | C | T | missense_variant | MODERATE | *AABR0702* | ENSRNOT00000032908.2 | 695 | c.695C>T | p.Pro232Leu | CC | CC | CC | TT | CC | CC | TT | CC |
| | | chr13:56263171 | T | A | missense_variant | MODERATE | *AABR0702* | ENSRNOT00000032908.2 | 929 | c.929T>A | p.Val310Glu | TT | TT | TT | AA | TT | TT | AA | TT |
| | | chr14:85271424 | G | A | missense_variant | MODERATE | *Ap1b1* | ENSRNOT00000057407.4 | 1850 | c.1822G>A | p.Ala608Thr | GG | GG | GG | GG | GG | GG | GG | AA |
| Fasting Glucose | chr14:86029588 | chr14:85271424 | G | A | missense_variant | MODERATE | *Ap1b1* | ENSRNOT00000089866.1 | 1987 | c.1822G>A | p.Ala608Thr | GG | GG | GG | GG | GG | GG | GG | AA |
| | | chr14:85756355 | A | G | missense_variant | MODERATE | *Xbp1* | ENSRNOT00000014044.6 | 397 | c.397A>G | p.Asn133Asp | AA | AA | AA | AA | AA | AA | AA | GG |
| | | chr14:85833986 | A | G | missense_variant | MODERATE | *Ankrd36* | ENSRNOT00000083756.1 | 1066 | c.857A>G | p.Glu286Gly | AA | AA | AA | AA | AA | AA | AA | GG |
| | | chr14:86071569 | T | C | missense_variant | MODERATE | *Polm* | ENSRNOT00000018319.6 | 1163 | c.916A>G | p.Met306Val | TT | TT | TT | TT | TT | TT | TT | CC |
| | | chr14:86071595 | G | A | missense_variant | MODERATE | *Polm* | ENSRNOT00000018319.6 | 1137 | c.890C>T | p.Ala297Val | GG | GG | GG | GG | GG | GG | GG | AA |
| | | chr14:86077302 | C | T | missense_variant | MODERATE | *Polm* | ENSRNOT00000018319.6 | 606 | c.359G>A | p.Arg120Gln | CC | CC | CC | CC | CC | CC | CC | TT |
| | | chr14:86077440 | T | C | missense_variant | MODERATE | *Polm* | ENSRNOT00000018319.6 | 468 | c.221A>G | p.Glu74Gly | TT | TT | TT | TT | TT | TT | TT | CC |
| | | chr14:86116112 | G | A | missense_variant | MODERATE | *Pold2* | ENSRNOT00000019288.4 | 263 | c.187C>T | p.Pro63Ser | GG | GG | GG | GG | GG | GG | GG | AA |
| | | chr14:86116112 | G | A | missense_variant | MODERATE | *Pold2* | ENSRNOT00000084633.1 | 252 | c.187C>T | p.Pro63Ser | GG | GG | GG | GG | GG | GG | GG | AA |
| | | chr14:86387570 | G | T | missense_variant | MODERATE | *Ddx56* | ENSRNOT00000089384.1 | 37 | c.17C>A | p.Ala6Glu | GG | GG | GG | GG | GG | GG | GG | TT |
| Body Length_NoTail | chr16:64014119 | chr16:64052952 | A | G | missense_variant | MODERATE | *Nrg1* | ENSRNOT00000014147.6 | 1422 | c.1078A>G | p.Ile360Val | GG | AA | GG | GG | GG | AA | GG | GG |
| | | chr16:64052952 | A | G | missense_variant | MODERATE | *Nrg1* | ENSRNOT00000013991.8 | 1876 | c.1255A>G | p.Ile419Val | GG | AA | GG | GG | GG | AA | GG | GG |
| | | chr16:64052952 | A | G | missense_variant | MODERATE | *Nrg1* | ENSRNOT00000014268.8 | 1500 | c.1156A>G | p.Ile386Val | GG | AA | GG | GG | GG | AA | GG | GG |
| | | chr16:64052952 | A | G | missense_variant | MODERATE | *Nrg1* | ENSRNOT00000081522.1 | 1891 | c.1270A>G | p.Ile424Val | GG | AA | GG | GG | GG | AA | GG | GG |
| | | chr16:64052952 | A | G | missense_variant | MODERATE | *Nrg1* | ENSRNOT00000082355.1 | 1876 | c.1255A>G | p.Ile419Val | GG | AA | GG | GG | GG | AA | GG | GG |
| | | chr16:64052952 | A | G | missense_variant | MODERATE | *Nrg1* | ENSRNOT00000058727.4 | 1446 | c.1102A>G | p.Ile368Val | GG | AA | GG | GG | GG | AA | GG | GG |
| | | chr16:64052952 | A | G | missense_variant | MODERATE | *Nrg1* | ENSRNOT00000038549.7 | 1320 | c.976A>G | p.Ile326Val | GG | AA | GG | GG | GG | AA | GG | GG |
| RetroFat | chr1:160530456 | chr1:162354117 | G | A | missense_variant | MODERATE | *Alg8* | ENSRNOT00000016478.6 | 891 | c.869G>A | p.Ser290Asn | AA | GG | GG | GG | GG | GG | AA | AA |
| | | chr1:162361693 | G | A | missense_variant | MODERATE | *Alg8* | ENSRNOT00000016478.6 | 1548 | c.1526G>A | p.Arg509Lys | GG | GG | GG | GG | GG | GG | AA | GG |
| BMI_Tail | chr1:282049439 | chr1:281755911 | C | T | start_lost | HIGH | *Prlhr* | ENSRNOT00000013170.4 | 23 | c.3G>A | p.Met1? | CC | CC | TT | CC | CC | CC | CC | TT |
| Body Weight | chr1:281756885 | chr1:281755911 | C | T | start_lost | HIGH | *Prlhr* | ENSRNOT00000013170.4 | 23 | c.3G>A | p.Met1? | CC | CC | TT | CC | CC | CC | CC | TT |
| ParaFat | chr1:280924549 | chr1:281755911 | C | T | start_lost | HIGH | *Prlhr* | ENSRNOT00000013170.4 | 23 | c.3G>A | p.Met1? | CC | CC | TT | CC | CC | CC | CC | TT |
| RetroFat | chr1:281777218 | chr1:281755911 | C | T | start_lost | HIGH | *Prlhr* | ENSRNOT00000013170.4 | 23 | c.3G>A | p.Met1? | CC | CC | TT | CC | CC | CC | CC | TT |
| EpiFat | chr1:281802657 | chr1:281755911 | C | T | start_lost | HIGH | *Prlhr* | ENSRNOT00000013170.4 | 23 | c.3G>A | p.Met1? | CC | CC | TT | CC | CC | CC | CC | TT |
| Body Weight | chr2:65816485 | chr2:62875814 | C | G | missense_variant | MODERATE | *RGD13065* | ENSRNOT00000039234.3 | 1109 | c.1004G>C | p.Ser335Thr | TT | CC | TT | TT | TT | TT | TT | TT |

| | | | | | | | | | | | | | | | | | | |
|---|---|---|---|---|---|---|---|---|---|---|---|---|---|---|---|---|---|---|
| Body Weight | chr2:[illegible] | chr2:62875821 | G | C | missense_variant | MODERATE | *RGD13065* | ENSRNOT00000039234.3 | 1102 | c.997C>G | p.Leu333Val | AA | GG | AA | AA | AA | AA | AA | AA |
| RetroFat | chr3:137537161 | chr3:136742468 | T | G | missense_variant | MODERATE | *Kif16b* | ENSRNOT00000036273.4 | 2319 | c.2263A>C | p.Lys755Gln | GG | TT | TT | TT | GG | GG | TT | TT |
| | | chr3:136742468 | T | G | missense_variant | MODERATE | *Kif16b* | ENSRNOT00000083061.1 | 2474 | c.2260A>C | p.Lys754Gln | GG | TT | TT | TT | GG | GG | TT | TT |
| | | chr3:138610633 | G | A | missense_variant | MODERATE | *Zfp133* | ENSRNOT00000031623.3 | 375 | c.242G>A | p.Arg81Gln | GG | GG | GG | GG | GG | GG | GG | AA |
| | | chr3:138615104 | T | A | missense_variant | MODERATE | *Zfp133* | ENSRNOT00000031623.3 | 1253 | c.1120T>A | p.Phe374Ile | TT | TT | TT | TT | TT | TT | TT | AA |
| RetroFat | chr3:95389621 | chr3:92494088 | A | C | missense_variant | MODERATE | *Pamr1* | ENSRNOT00000064282.2 | 1026 | c.905A>C | p.Glu302Ala | CC | AA | CC | CC | AA | CC | AA | AA |
| | | chr3:92501374 | A | G | missense_variant | MODERATE | *Pamr1* | ENSRNOT00000064282.2 | 2096 | c.1975A>G | p.Asn659Asp | AA | AA | AA | AA | AA | AA | GG | AA |
| | | chr3:92910764 | A | G | missense_variant | MODERATE | *Pdhx* | ENSRNOT00000009552.5 | 1196 | c.1156T>C | p.Tyr386His | AA | AA | AA | AA | GG | AA | GG | AA |
| | | chr3:92924011 | G | C | missense_variant | MODERATE | *Pdhx* | ENSRNOT00000088242.1 | 842 | c.748C>G | p.Pro250Ala | GG | GG | GG | GG | CC | GG | CC | GG |
| | | chr3:92924011 | G | C | missense_variant | MODERATE | *Pdhx* | ENSRNOT00000009552.5 | 746 | c.706C>G | p.Pro236Ala | GG | GG | GG | GG | CC | GG | CC | GG |
| | | chr3:92947902 | C | T | missense_variant | MODERATE | *Pdhx* | ENSRNOT00000088242.1 | 300 | c.206G>A | p.Arg69Gln | CC | CC | CC | CC | TT | CC | TT | CC |
| | | chr3:94018570 | G | A | missense_variant | MODERATE | *Cd59* | ENSRNOT00000067085.3 | 102 | c.11G>A | p.Arg4Gln | GG | GG | GG | GG | AA | GG | AA | GG |
| | | chr3:94353600 | C | A | missense_variant | MODERATE | *Hipk3* | ENSRNOT00000089554.1 | 3343 | c.3210G>T | p.Leu1070Phe | CC | CC | CC | CC | AA | CC | AA | CC |
| | | chr3:94353600 | C | A | missense_variant | MODERATE | *Hipk3* | ENSRNOT00000015775.3 | 3614 | c.3144G>T | p.Leu1048Phe | CC | CC | CC | CC | AA | CC | AA | CC |
| | | chr3:94982464 | C | T | missense_variant | MODERATE | *Ccdc73* | ENSRNOT00000041362.5 | 2435 | c.2219C>T | p.Pro740Leu | CC | CC | CC | CC | TT | CC | TT | CC |
| | | chr3:97349458 | G | A | missense_variant | MODERATE | *Dcdc5* | ENSRNOT00000041494.4 | 359 | c.359G>A | p.Arg120His | GG | GG | GG | GG | GG | GG | AA | GG |
| | | chr3:97396118 | G | A | missense_variant | MODERATE | *Dcdc5* | ENSRNOT00000041494.4 | 1307 | c.1307G>A | p.Ser436Asn | GG | GG | GG | GG | GG | GG | AA | GG |
| | | chr3:97396118 | G | A | missense_variant | MODERATE | *Dcdc5* | ENSRNOT00000089524.1 | 1174 | c.1043G>A | p.Ser348Asn | GG | GG | GG | GG | GG | GG | AA | GG |
| | | chr3:97398578 | A | C | missense_variant | MODERATE | *Dcdc5* | ENSRNOT00000041494.4 | 1500 | c.1500A>C | p.Glu500Asp | AA | AA | AA | AA | AA | AA | CC | AA |
| | | chr3:97398578 | A | C | missense_variant | MODERATE | *Dcdc5* | ENSRNOT00000089524.1 | 1367 | c.1236A>C | p.Glu412Asp | AA | AA | AA | AA | AA | AA | CC | AA |
| Body Weight | chr5:50933779 | chr5:50066002 | G | A | missense_variant | MODERATE | *Orc3* | ENSRNOT00000011085.4 | 250 | c.250C>T | p.Leu84Phe | AA | GG | GG | AA | AA | GG | AA | GG |
| Body Length_Tail | chr6:137745191 | chr6:137323809 | G | A | missense_variant | MODERATE | *Pld4* | ENSRNOT00000029017.3 | 97 | c.77G>A | p.Arg26Lys | AA | GG | GG | GG | GG | AA | GG | AA |
| | | chr6:137326285 | T | C | missense_variant | MODERATE | *Pld4* | ENSRNOT00000029017.3 | 331 | c.311T>C | p.Phe104Ser | CC | TT | TT | TT | TT | CC | TT | CC |
| | | chr6:137337270 | G | A | missense_variant | MODERATE | *Ahnak2* | ENSRNOT00000039631.4 | 15980 | c.15785C>T | p.Pro5262Leu | AA | GG | GG | GG | GG | AA | GG | AA |
| | | chr6:137339352 | A | G | missense_variant | MODERATE | *Ahnak2* | ENSRNOT00000039631.4 | 13898 | c.13703T>C | p.Val4568Ala | GG | AA | AA | AA | AA | GG | AA | GG |
| | | chr6:137347877 | A | T | missense_variant | MODERATE | *Ahnak2* | ENSRNOT00000039631.4 | 5373 | c.5178T>A | p.His1726Gln | TT | AA | AA | AA | AA | TT | AA | TT |
| | | chr6:137347903 | G | A | missense_variant | MODERATE | *Ahnak2* | ENSRNOT00000039631.4 | 5347 | c.5152C>T | p.Pro1718Ser | AA | GG | GG | GG | GG | AA | GG | AA |
| | | chr6:137348049 | A | G | missense_variant | MODERATE | *Ahnak2* | ENSRNOT00000039631.4 | 5201 | c.5006T>C | p.Val1669Ala | GG | AA | AA | AA | AA | GG | AA | GG |
| | | chr6:137348399 | C | T | missense_variant | MODERATE | *Ahnak2* | ENSRNOT00000039631.4 | 4851 | c.4656G>A | p.Met1552Ile | TT | CC | CC | CC | CC | TT | CC | TT |
| | | chr6:137348644 | T | C | missense_variant | MODERATE | *Ahnak2* | ENSRNOT00000039631.4 | 4606 | c.4411A>G | p.Asn1471Asp | CC | TT | TT | TT | TT | CC | TT | CC |
| | | chr6:137348665 | A | G | missense_variant | MODERATE | *Ahnak2* | ENSRNOT00000039631.4 | 4585 | c.4390T>C | p.Trp1464Arg | GG | AA | AA | AA | AA | GG | AA | GG |
| | | chr6:137876105 | T | C | missense_variant | MODERATE | *Pacs2* | ENSRNOT00000056880.5 | 1111 | c.1067T>C | p.Met356Thr | CC | TT | TT | TT | TT | TT | TT | TT |
| | | chr6:137885119 | T | C | missense_variant | MODERATE | *Pacs2* | ENSRNOT00000056880.5 | 2278 | c.2234T>C | p.Val745Ala | CC | TT | TT | TT | TT | CC | TT | TT |
| | | chr6:137970275 | G | C | missense_variant | MODERATE | *LOC69042* | ENSRNOT00000029223.6 | 574 | c.379G>C | p.Val127Leu | CC | GG | GG | GG | GG | CC | GG | GG |
| | | chr6:138068384 | T | C | missense_variant | MODERATE | *Ighm* | ENSRNOT00000081908.1 | 265 | c.220A>G | p.Ile74Val | CC | TT | TT | TT | TT | CC | TT | TT |
| EpiFat | chr6:26266960 | chr6:23340330 | A | T | missense_variant | MODERATE | *RGD13049* | ENSRNOT00000011832.7 | 2760 | c.2520A>T | p.Glu840Asp | TT | AA | AA | AA | AA | AA | AA | TT |
| | | chr6:23341333 | A | G | missense_variant | MODERATE | *RGD13049* | ENSRNOT00000011832.7 | 3763 | c.3523A>G | p.Thr1175Ala | GG | AA | AA | AA | AA | AA | AA | GG |
| | | chr6:26099033 | G | A | missense_variant | MODERATE | *Rbks* | ENSRNOT00000006452.6 | 1039 | c.787G>A | p.Val263Met | AA | GG | GG | GG | GG | GG | GG | AA |
| | | chr6:26135860 | A | G | missense_variant | MODERATE | *Mrpl33* | ENSRNOT00000034712.4 | 94 | c.94T>C | p.Tyr32His | GG | AA | AA | AA | AA | AA | AA | GG |
| | | chr6:26382623 | G | A | missense_variant | MODERATE | *Gckr* | ENSRNOT00000073228.2 | 622 | c.445C>T | p.Arg149Cys | GG | GG | GG | GG | GG | GG | GG | AA |
| | | chr6:26382623 | G | A | missense_variant | MODERATE | *Gckr* | ENSRNOT00000073228.2 | 622 | c.445C>T | p.Arg149Cys | GG | GG | GG | GG | GG | GG | GG | AA |
| | | chr6:26786379 | A | G | missense_variant | MODERATE | *Preb* | ENSRNOT00000009565.6 | 1051 | c.881A>G | p.Gln294Arg | AA | AA | AA | AA | AA | AA | AA | GG |
| | | chr6:26786379 | A | G | missense_variant | MODERATE | *Preb* | ENSRNOT00000009565.6 | 1051 | c.881A>G | p.Gln294Arg | AA | AA | AA | AA | AA | AA | AA | GG |
| | | chr6:27428501 | G | A | missense_variant | MODERATE | *Drc1* | ENSRNOT00000036815.5 | 1856 | c.1811C>T | p.Thr604Ile | GG | GG | GG | GG | GG | GG | GG | AA |
| | | chr6:27428501 | G | A | missense_variant | MODERATE | *Drc1* | ENSRNOT00000036815.5 | 1856 | c.1811C>T | p.Thr604Ile | GG | GG | GG | GG | GG | GG | GG | AA |
| RetroFat | chr6:28148338 | chr6:28004664 | A | G | missense_variant | MODERATE | *Dtnb* | ENSRNOT00000077830.1 | 421 | c.308A>G | p.Asn103Ser | AA | AA | AA | AA | AA | AA | AA | GG |
| | | chr6:28004664 | A | G | missense_variant | MODERATE | *Dtnb* | ENSRNOT00000060810.2 | 594 | c.308A>G | p.Asn103Ser | AA | AA | AA | AA | AA | AA | AA | GG |
| | | chr6:28004664 | A | G | missense_variant | MODERATE | *Dtnb* | ENSRNOT00000077830.1 | 421 | c.308A>G | p.Asn103Ser | AA | AA | AA | AA | AA | AA | AA | GG |
| | | chr6:28004664 | A | G | missense_variant | MODERATE | *Dtnb* | ENSRNOT00000060810.2 | 594 | c.308A>G | p.Asn103Ser | AA | AA | AA | AA | AA | AA | AA | GG |
| | | chr6:28572363 | T | C | missense_variant | MODERATE | *Adcy3* | ENSRNOT00000005389.6 | 585 | c.362T>C | p.Leu121Pro | TT | TT | TT | TT | TT | TT | TT | CC |
| Body Weight | chr7:36497588 | chr7:34310284 | T | G | missense_variant | MODERATE | *Lta4h* | ENSRNOT00000005930.4 | 1407 | c.1314T>G | p.Asp438Glu | TT | TT | GG | TT | TT | TT | TT | TT |
| Body Length_NoTail | chr7:36517726 | chr7:34310284 | T | G | missense_variant | MODERATE | *Lta4h* | ENSRNOT00000005930.4 | 1407 | c.1314T>G | p.Asp438Glu | TT | TT | GG | TT | TT | TT | TT | TT |
| Body Length_Tail | chr7:36517726 | chr7:34310284 | T | G | missense_variant | MODERATE | *Lta4h* | ENSRNOT00000005930.4 | 1407 | c.1314T>G | p.Asp438Glu | TT | TT | GG | TT | TT | TT | TT | TT |
| | | chr7:34889309 | T | C | missense_variant | MODERATE | *Vezt* | ENSRNOT00000030015.4 | 2061 | c.2002A>G | p.Met668Val | TT | TT | CC | TT | CC | CC | CC | TT |
| | | chr7:34889309 | T | C | missense_variant | MODERATE | *Vezt* | ENSRNOT00000030015.4 | 2061 | c.2002A>G | p.Met668Val | TT | TT | CC | TT | CC | CC | CC | TT |
| | | chr8:116614891 | G | C | missense_variant | MODERATE | *Rbm6* | ENSRNOT00000024939.5 | 1333 | c.1093C>G | p.Gln365Glu | CC | GG | GG | GG | GG | GG | GG | GG |
| | | chr8:117289662 | C | A | missense_variant | MODERATE | *Usp19* | ENSRNOT00000085038.1 | 4102 | c.3803C>A | p.Ala1268Asp | CC | CC | CC | AA | CC | CC | CC | CC |
| | | chr8:117289662 | C | A | missense_variant | MODERATE | *Usp19* | ENSRNOT00000074772.2 | 3797 | c.3797C>A | p.Ala1266Asp | CC | CC | CC | AA | CC | CC | CC | CC |
| | | chr8:117661846 | C | T | missense_variant | MODERATE | *Tmem89* | ENSRNOT00000049240.5 | 496 | c.470C>T | p.Ala157Val | CC | CC | CC | TT | CC | TT | TT | TT |

| | | | | | | | | | | | | | | | | | | | |
|---|---|---|---|---|---|---|---|---|---|---|---|---|---|---|---|---|---|---|---|
| Body Length_Tail | chr8:118711320 | chr8:117706643 | A | G | missense_variant | MODERATE | *Col7a1* | ENSRNOT00000027994.6 | 4258 | c.4258A>G | p.Ser1420Gly | GG | AA | AA | AA | AA | AA | AA | AA |
| | | chr8:117714681 | G | A | missense_variant | MODERATE | *Col7a1* | ENSRNOT00000027994.6 | 6034 | c.6034G>A | p.Gly2012Ser | GG | GG | GG | AA | GG | AA | AA | AA |
| | | chr8:117796955 | G | A | missense_variant | MODERATE | *Trex1* | ENSRNOT00000033719.5 | 192 | c.167C>T | p.Pro56Leu | AA | GG | GG | GG | GG | GG | GG | GG |
| | | chr8:117931377 | T | C | missense_variant | MODERATE | *Camp* | ENSRNOT00000028130.7 | 496 | c.344A>G | p.Gln115Arg | CC | TT | CC | TT | CC | TT | TT | TT |
| | | chr8:118119545 | G | A | missense_variant | MODERATE | *Map4* | ENSRNOT00000056161.4 | 2882 | c.2864G>A | p.Arg955Gln | GG | GG | GG | AA | GG | AA | AA | AA |
| | | chr8:118121377 | A | G | missense_variant | MODERATE | *Map4* | ENSRNOT00000056161.4 | 4714 | c.4696A>G | p.Asn1566Asp | AA | AA | AA | GG | AA | GG | GG | GG |
| | | chr8:118378322 | C | T | missense_variant | MODERATE | *RGD15637* | ENSRNOT00000047247.4 | 139 | c.139G>A | p.Ala47Thr | TT | CC | CC | CC | CC | CC | CC | CC |
| | | chr8:118664419 | C | T | stop_gained | HIGH | *Ngp* | ENSRNOT00000029755.2 | 329 | c.316C>T | p.Gln106* | TT | CC | CC | CC | CC | CC | CC | CC |
| | | chr8:118664420 | A | C | missense_variant | MODERATE | *Ngp* | ENSRNOT00000029755.2 | 330 | c.317A>C | p.Gln106Pro | CC | AA | AA | AA | AA | AA | AA | AA |
| | | chr8:118822162 | T | C | missense_variant | MODERATE | *Setd2* | ENSRNOT00000087154.1 | 716 | c.485T>C | p.Val162Ala | TT | TT | TT | CC | TT | CC | CC | CC |
| | | chr8:118822165 | C | T | missense_variant | MODERATE | *Setd2* | ENSRNOT00000087154.1 | 719 | c.488C>T | p.Ala163Val | CC | CC | CC | TT | CC | TT | TT | TT |
| | | chr8:118822336 | C | T | missense_variant | MODERATE | *Setd2* | ENSRNOT00000087154.1 | 890 | c.659C>T | p.Ala220Val | CC | CC | CC | TT | CC | TT | TT | TT |
| | | chr8:118824592 | A | G | missense_variant | MODERATE | *Setd2* | ENSRNOT00000028409.6 | 2656 | c.2228A>G | p.Glu743Gly | GG | AA | AA | AA | AA | AA | AA | AA |
| | | chr8:118824592 | A | G | missense_variant | MODERATE | *Setd2* | ENSRNOT00000087154.1 | 3146 | c.2915A>G | p.Glu972Gly | GG | AA | AA | AA | AA | AA | AA | AA |
| | | chr8:118825098 | G | C | missense_variant | MODERATE | *Setd2* | ENSRNOT00000028409.6 | 3162 | c.2734G>C | p.Val912Leu | GG | GG | GG | CC | GG | CC | CC | CC |
| | | chr8:118825098 | G | C | missense_variant | MODERATE | *Setd2* | ENSRNOT00000087154.1 | 3652 | c.3421G>C | p.Val1141Leu | GG | GG | GG | CC | GG | CC | CC | CC |
| | | chr8:118825125 | A | G | missense_variant | MODERATE | *Setd2* | ENSRNOT00000028409.6 | 3189 | c.2761A>G | p.Ile921Val | AA | AA | AA | GG | AA | GG | GG | GG |
| | | chr8:118825125 | A | G | missense_variant | MODERATE | *Setd2* | ENSRNOT00000087154.1 | 3679 | c.3448A>G | p.Ile1150Val | AA | AA | AA | GG | AA | GG | GG | GG |
| | | chr8:118825138 | T | C | missense_variant | MODERATE | *Setd2* | ENSRNOT00000028409.6 | 3202 | c.2774T>C | p.Met925Thr | TT | TT | TT | CC | TT | CC | CC | CC |
| | | chr8:118825138 | T | C | missense_variant | MODERATE | *Setd2* | ENSRNOT00000087154.1 | 3692 | c.3461T>C | p.Met1154Thr | TT | TT | TT | CC | TT | CC | CC | CC |
| | | chr8:118890799 | T | C | missense_variant | MODERATE | *Nradd* | ENSRNOT00000028416.4 | 801 | c.476A>G | p.Gln159Arg | TT | TT | TT | CC | TT | CC | CC | CC |
| | | chr8:118897582 | C | T | missense_variant | MODERATE | *Nbeal2* | ENSRNOT00000056130.5 | 6917 | c.6719G>A | p.Arg2240Gln | TT | CC | TT | CC | TT | CC | CC | CC |
| | | chr8:118901536 | T | C | missense_variant | MODERATE | *Nbeal2* | ENSRNOT00000056130.5 | 5161 | c.4963A>G | p.Met1655Val | CC | TT | CC | TT | CC | TT | TT | TT |
| | | chr8:118903111 | G | A | missense_variant | MODERATE | *Nbeal2* | ENSRNOT00000056130.5 | 4181 | c.3983C>T | p.Pro1328Leu | AA | GG | AA | GG | AA | GG | GG | GG |
| | | chr8:119084015 | G | A | missense_variant | MODERATE | *Prss44* | ENSRNOT00000051947.5 | 116 | c.82G>A | p.Val28Ile | AA | GG | AA | GG | AA | GG | GG | GG |
| | | chr8:119084015 | G | A | missense_variant | MODERATE | *Prss44* | ENSRNOT00000048655.5 | 95 | c.82G>A | p.Val28Ile | AA | GG | AA | GG | AA | GG | GG | GG |
| | | chr8:119115774 | G | A | missense_variant | MODERATE | *Prss45* | ENSRNOT00000043249.2 | 221 | c.173G>A | p.Arg58His | AA | GG | GG | GG | GG | GG | GG | GG |
| | | chr8:119139251 | A | G | missense_variant | MODERATE | *Prss50* | ENSRNOT00000056114.3 | 676 | c.676A>G | p.Lys226Glu | GG | AA | AA | AA | AA | AA | AA | AA |
| | | chr8:119160764 | C | A | missense_variant | MODERATE | *Als2cl* | ENSRNOT00000046745.4 | 34 | c.34C>A | p.Leu12Met | AA | CC | CC | CC | CC | CC | CC | CC |
| | | chr8:119236824 | T | A | missense_variant | MODERATE | *Lrrc2* | ENSRNOT00000043737.5 | 317 | c.251T>A | p.Val84Glu | TT | TT | AA | AA | AA | AA | AA | AA |
| | | chr8:119236824 | T | A | missense_variant | MODERATE | *Lrrc2* | ENSRNOT00000078439.1 | 435 | c.251T>A | p.Val84Glu | TT | TT | AA | AA | AA | AA | AA | AA |
| | | chr8:119236869 | A | G | missense_variant | MODERATE | *Lrrc2* | ENSRNOT00000043737.5 | 362 | c.296A>G | p.Asn99Ser | AA | AA | GG | GG | GG | GG | GG | GG |
| | | chr8:119236869 | A | G | missense_variant | MODERATE | *Lrrc2* | ENSRNOT00000078439.1 | 480 | c.296A>G | p.Asn99Ser | AA | AA | GG | GG | GG | GG | GG | GG |
| | | chr8:119640095 | A | G | missense_variant | MODERATE | *Trank1* | ENSRNOT00000028633.5 | 5963 | c.5963A>G | p.Lys1988Arg | GG | AA | AA | AA | AA | GG | GG | AA |
| | | chr8:119640688 | C | G | missense_variant | MODERATE | *Trank1* | ENSRNOT00000028633.5 | 6556 | c.6556C>G | p.Leu2186Val | GG | CC | CC | CC | CC | GG | GG | CC |
| | | chr8:119691654 | G | A | missense_variant&sp | MODERATE | *Dclk3* | ENSRNOT00000044742.5 | 267 | c.88G>A | p.Gly30Ser | AA | GG | GG | GG | GG | AA | AA | GG |
| Body Length_Tail | chr9:65078205 | chr9:64913080 | A | G | missense_variant | MODERATE | *Sgo2* | ENSRNOT00000029526.6 | 1024 | c.785A>G | p.Lys262Arg | GG | AA | GG | AA | GG | GG | GG | GG |
| | | chr9:64953533 | G | A | missense_variant | MODERATE | *Aox1* | ENSRNOT00000068633.4 | 376 | c.328G>A | p.Gly110Ser | GG | GG | GG | AA | GG | GG | GG | GG |
| | | chr9:65026169 | G | A | missense_variant | MODERATE | *Aox3* | ENSRNOT00000081146.1 | 535 | c.458G>A | p.Arg153His | GG | GG | GG | AA | GG | GG | GG | GG |
| | | chr9:65063124 | G | A | missense_variant | MODERATE | *Aox3* | ENSRNOT00000092906.1 | 2430 | c.1688G>A | p.Gly563Asp | GG | GG | GG | AA | GG | GG | GG | GG |
| | | chr9:65063124 | G | A | missense_variant | MODERATE | *Aox3* | ENSRNOT00000081146.1 | 2428 | c.2351G>A | p.Gly784Asp | GG | GG | GG | AA | GG | GG | GG | GG |

**Supplementary Figure S1**

Genetic and phenotypic correlation between adiposity traits, fasting glucose and tail length. Phenotypic correlations are depicted in the upper part, genetic – in the lower part of the matrix. Number inside squares show p-value > 0.05.

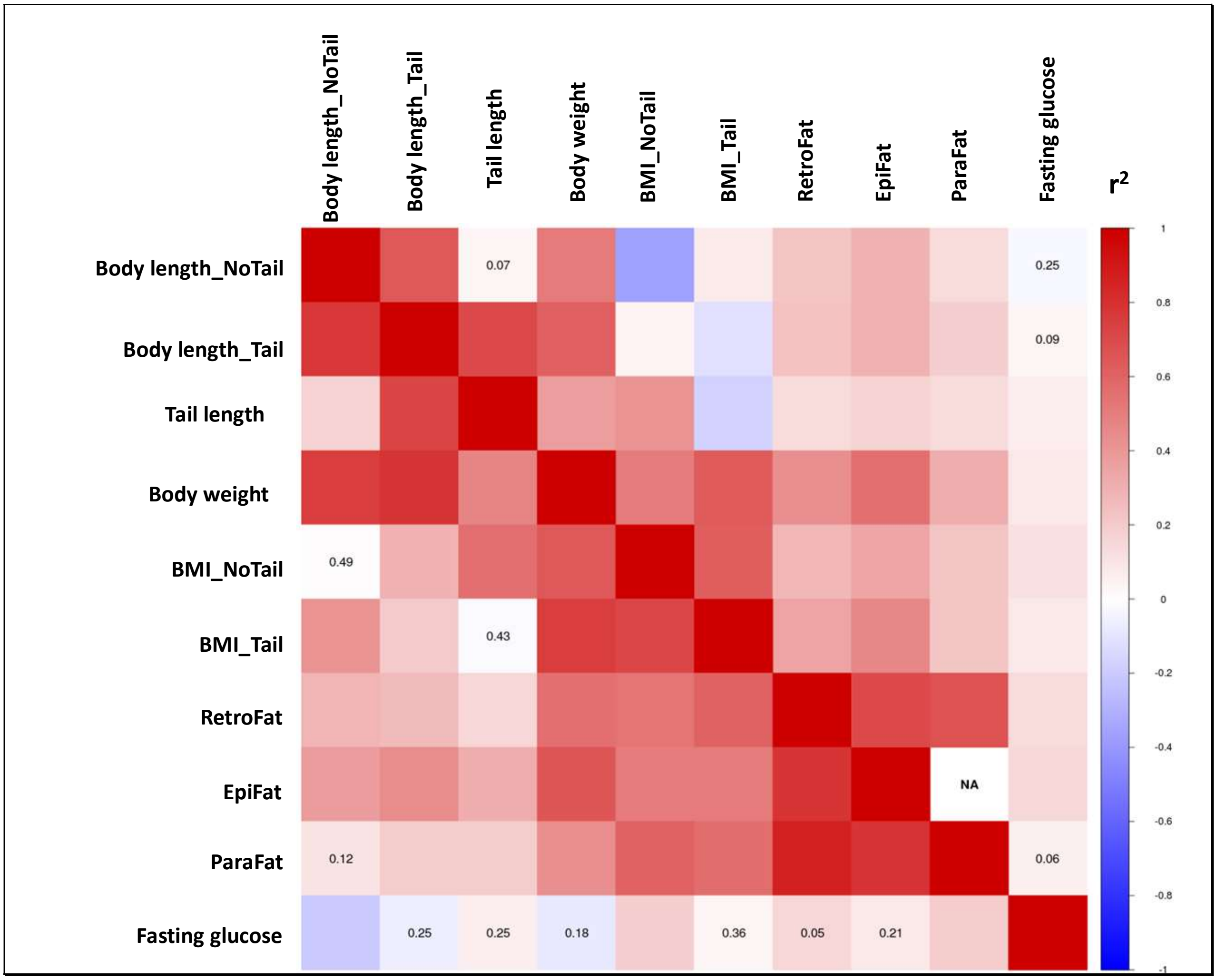

**Supplementary Figure S2**

**Manhattan plots. Genome-wide association results from the GWA analysis for 9 adiposity traits. The chromosomal distribution of all the P-values (-$\log_{10}$P values) is shown**

**Retroperitoneal fat weight**

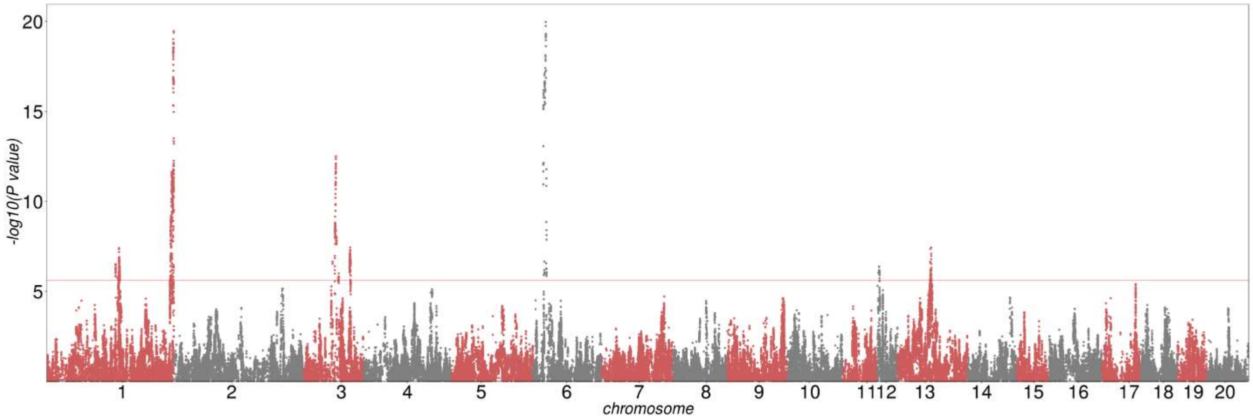


**Body weight**

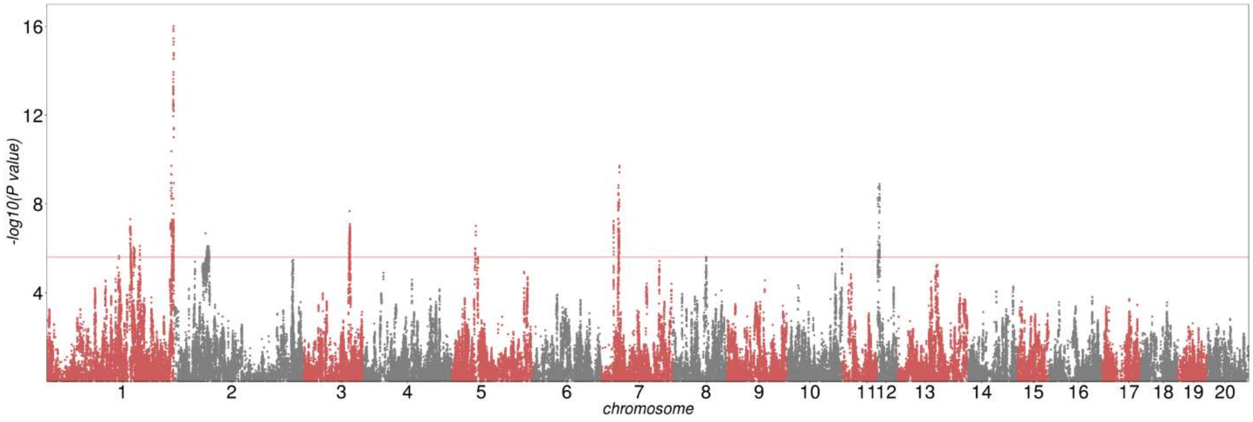

### Body length with tail

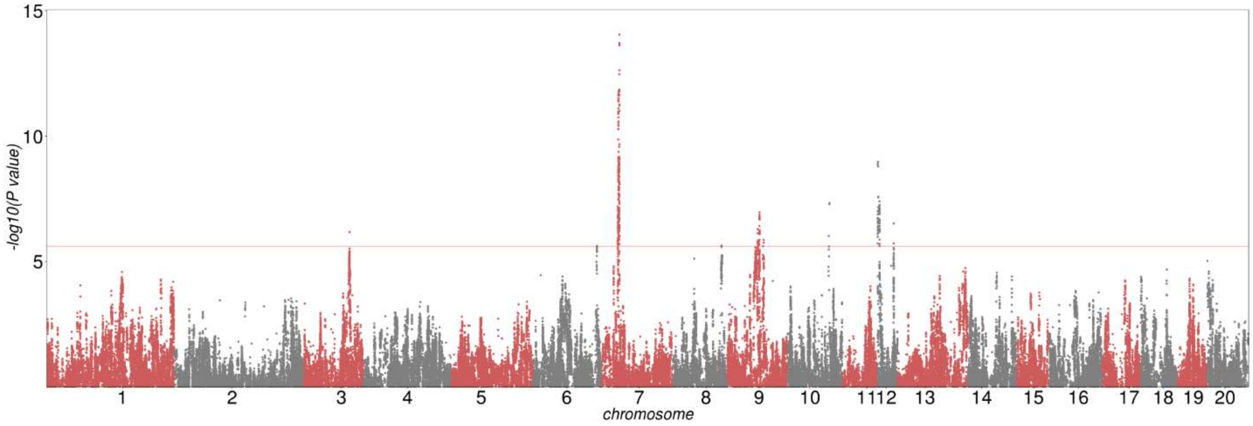


### Body length without tail

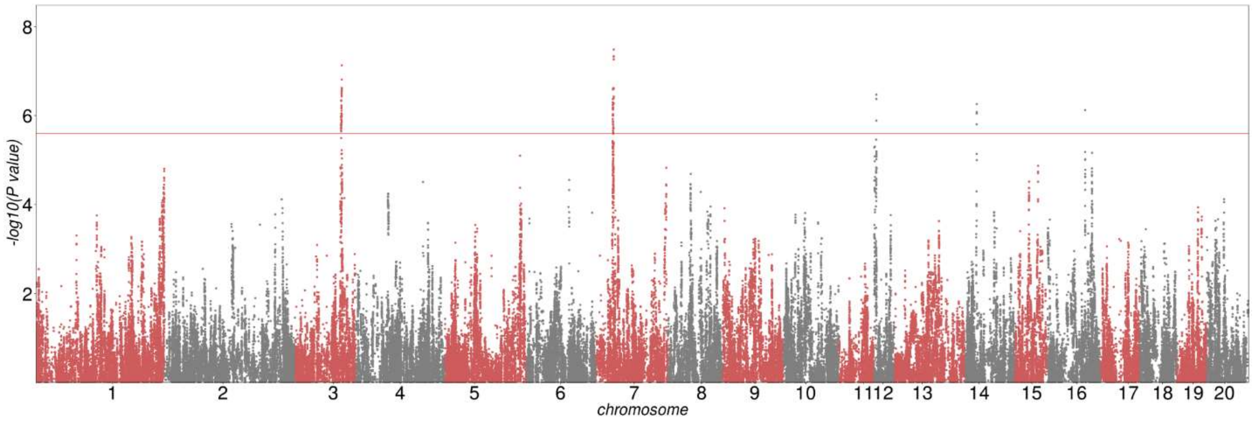

**BMI – body length with tail**

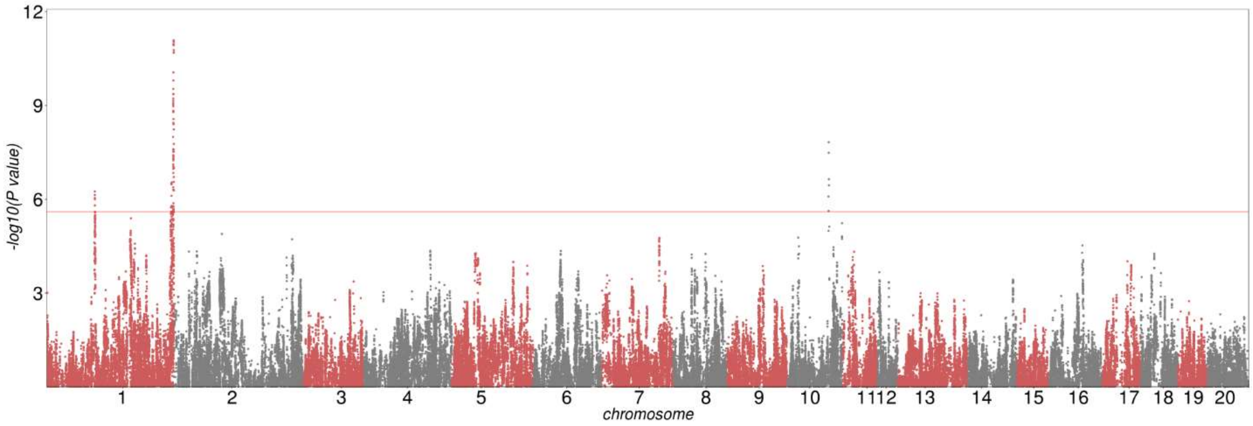


**BMI – body length without tail**

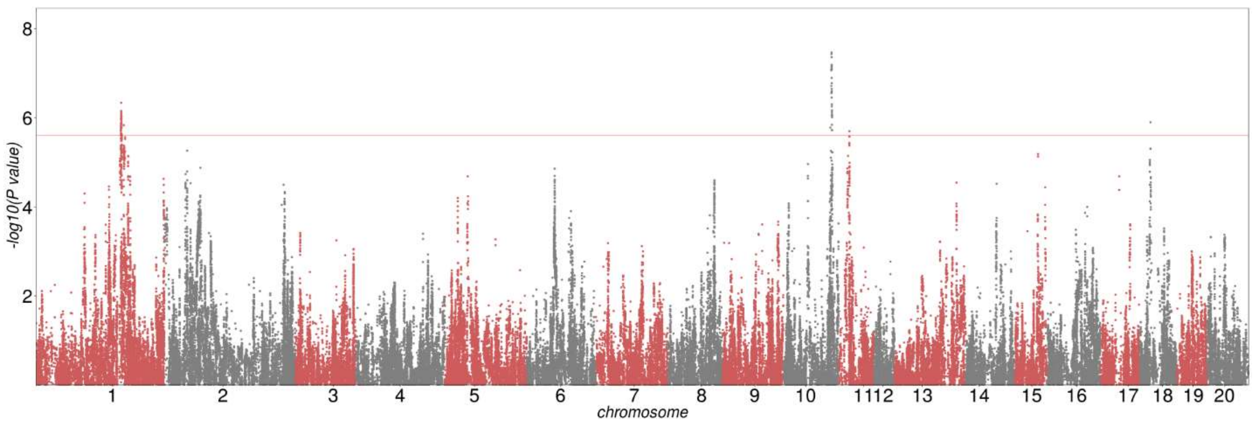

**Epididymis fat weight**

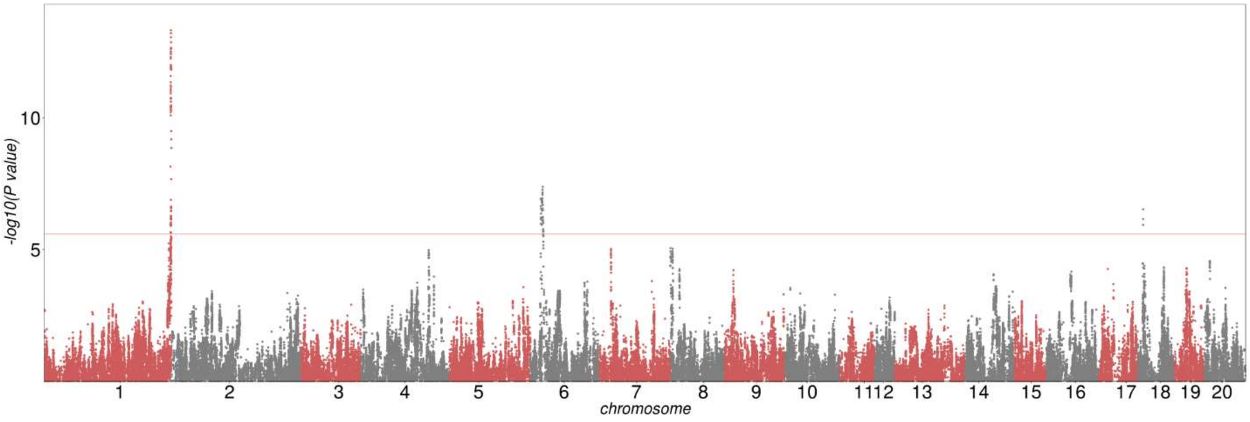


**Parametrial fat weight**

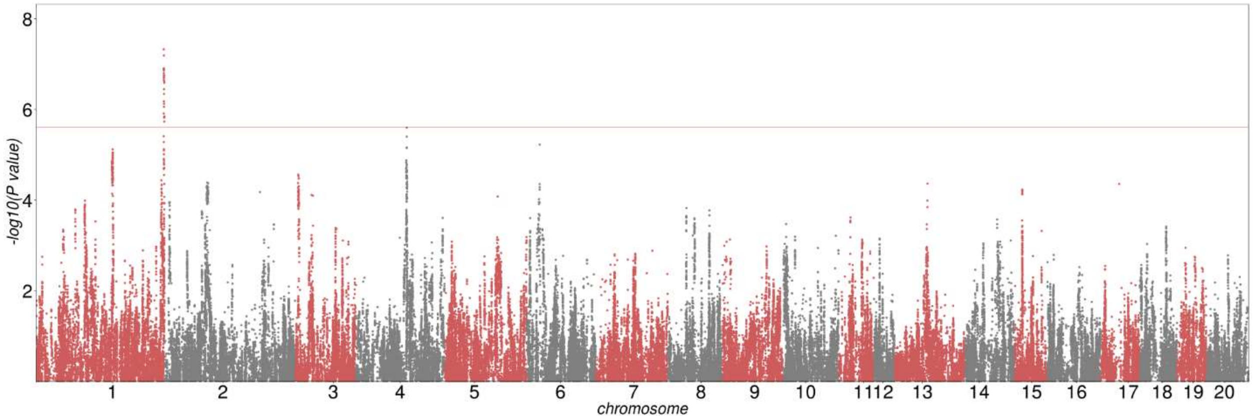

**Fasting glucose**

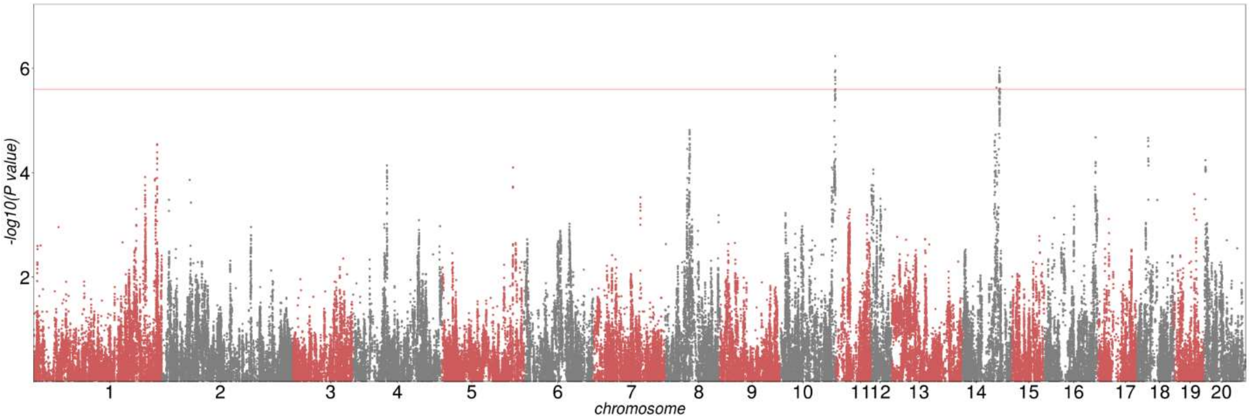


**Tail length**

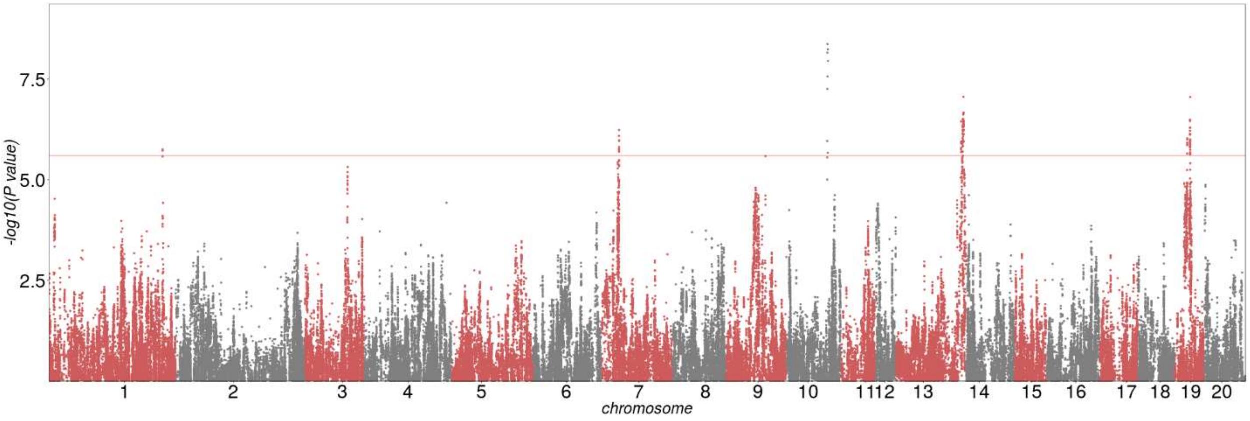

**Supplementary Figure S3.** 46 QTLs for 10 traits. Regional association plots show the vicinity of the top SNPs for each QTL. The SNPs with the lowest p-value (“top SNP) is shown in purple. Correlation of each SNP with the top SNP is shown in color. Credible set track shows the smallest set of SNPs accounting for 99% of the posterior probability (“credible set”). Genes in the region were annotated using Refseq annotation. The chromosome regions were chosen to optimally show LD structure of the region

BMI_Tail, Chr1:106,866,154

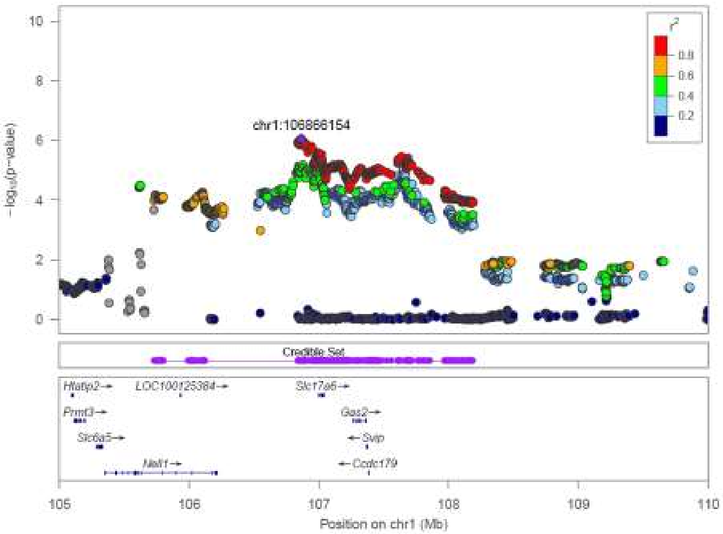

RetroFat, Chr1:160,530,456

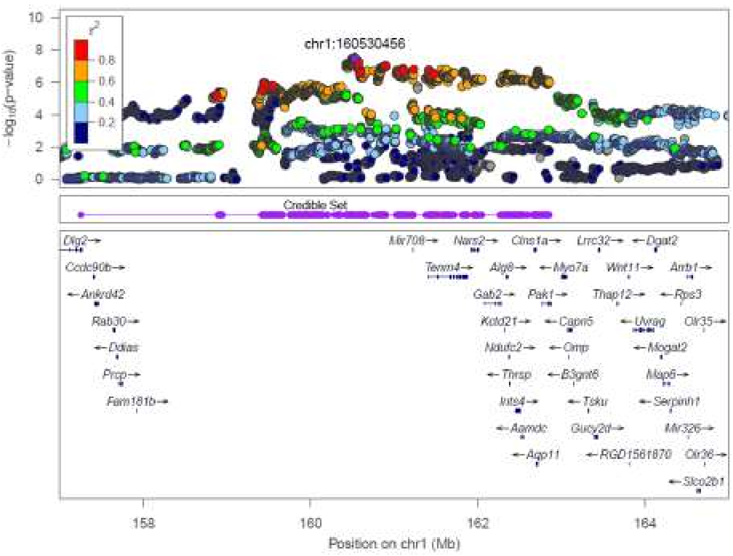


Body weight, Chr1:185,730,317

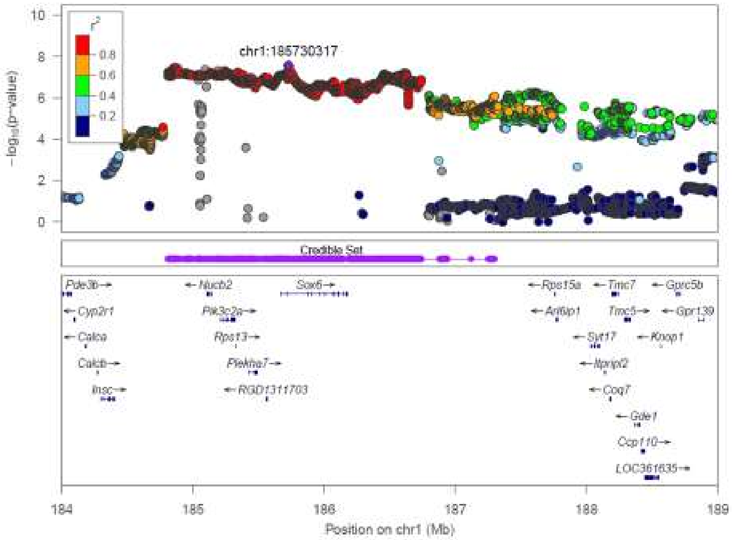

BMI_NoTail, Chr1:187,300,775

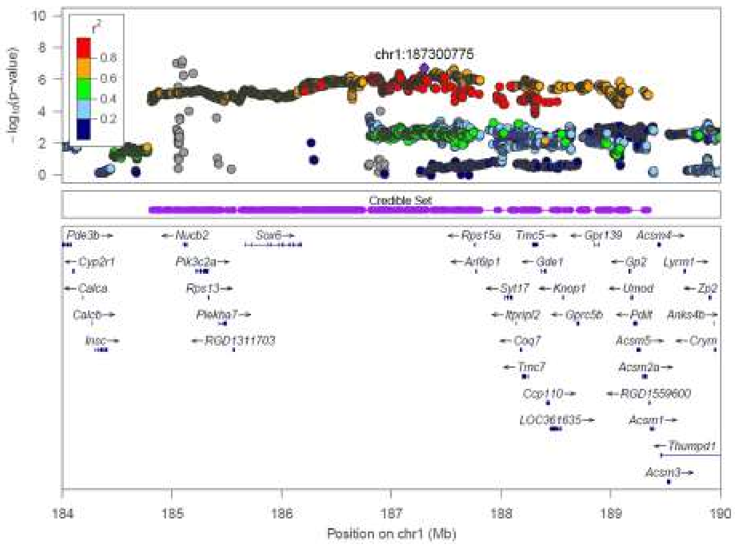

TL, Chr1:253,524,003

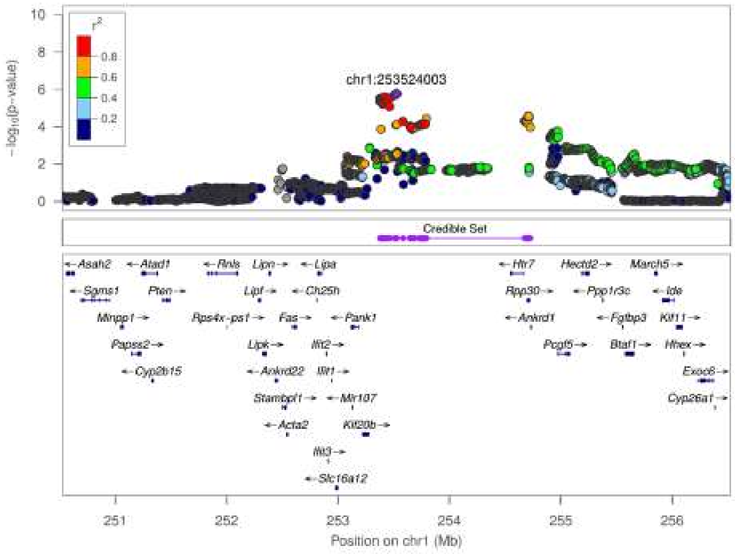

ParaFat, Chr1:280,924,549

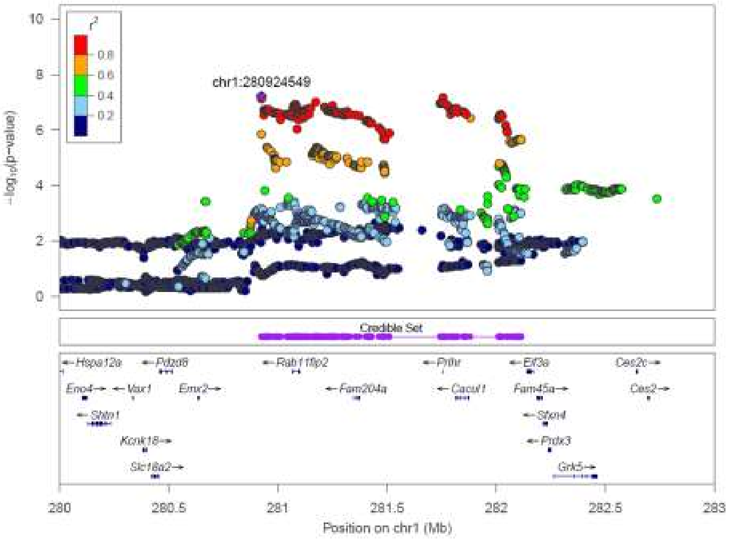


Body weight, Chr1:281,756,885

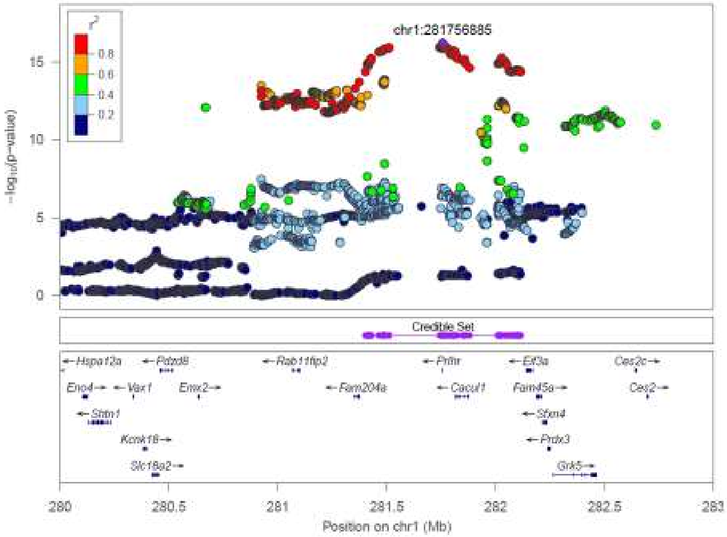

RetroFat, Chr1:281,777,218

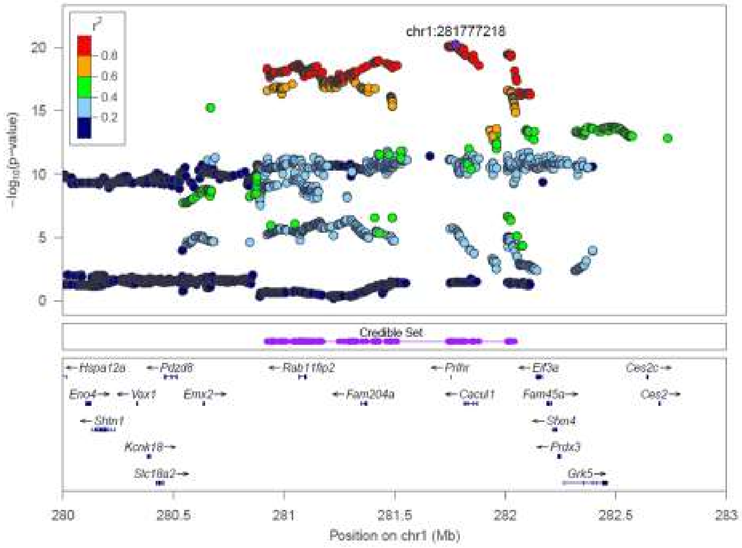


EpiFat, Chr1:281,802,657

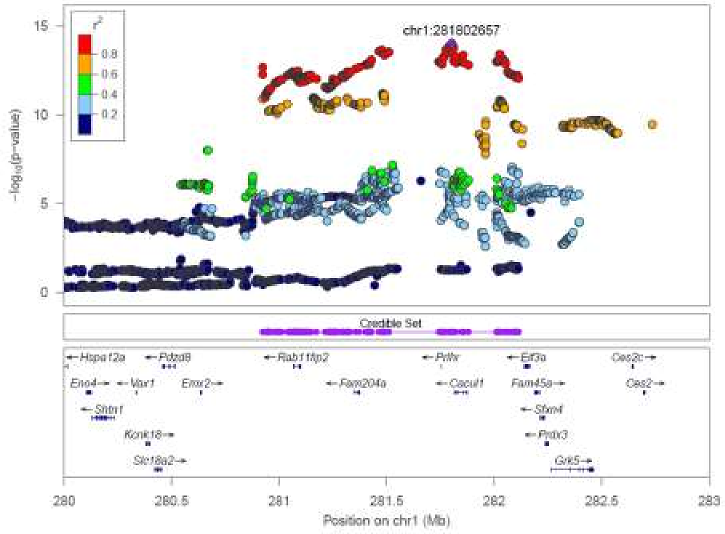

BMI_Tail, Chr1:282,049,439

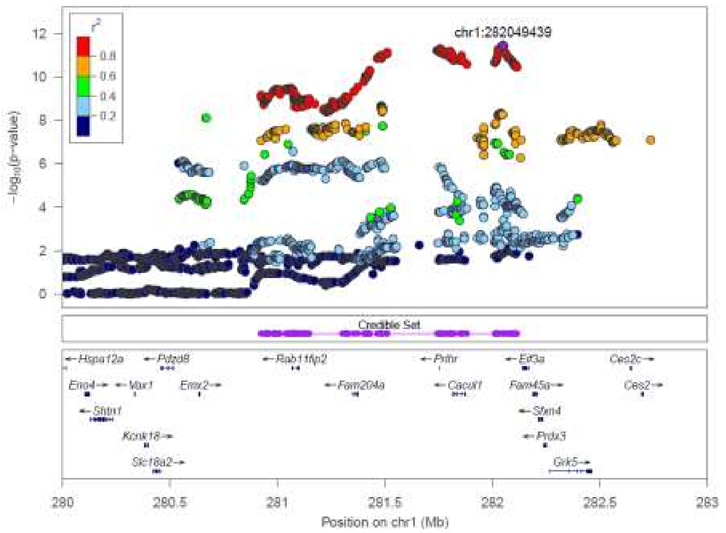


Body weight, Chr2:65,816,485

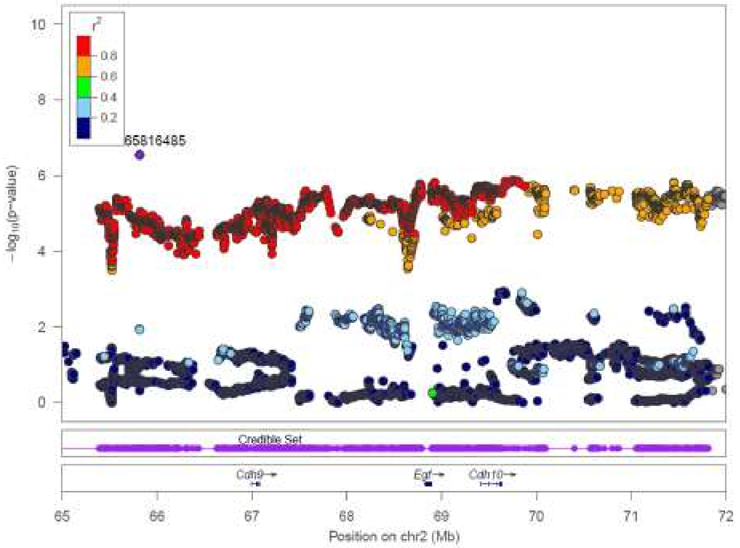

RetroFat, Chr3:95,389,621

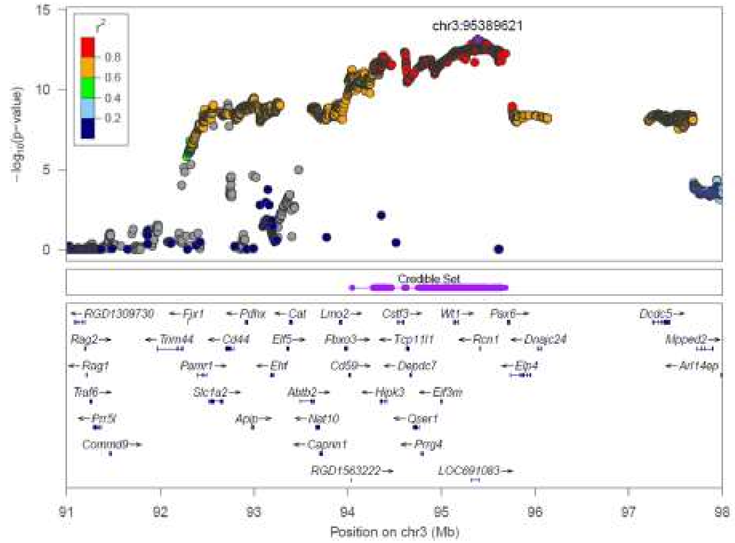

Body weight, Chr3:136,021,511

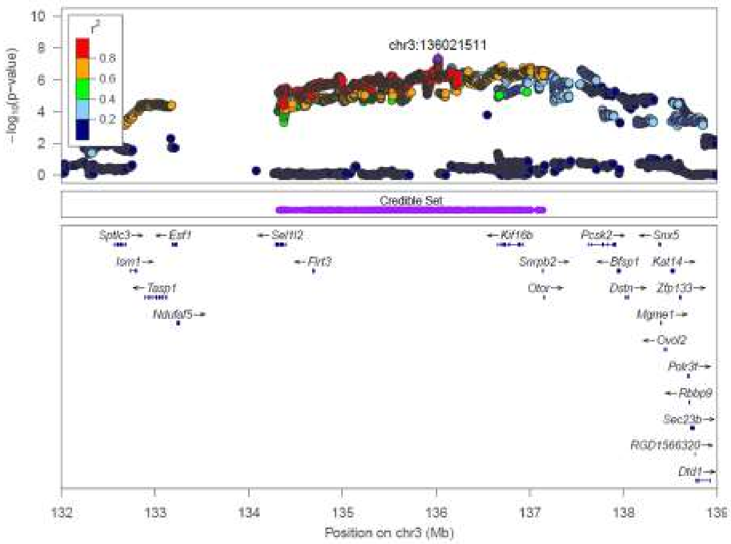

Body Length_Tail, Chr3:136,021,511

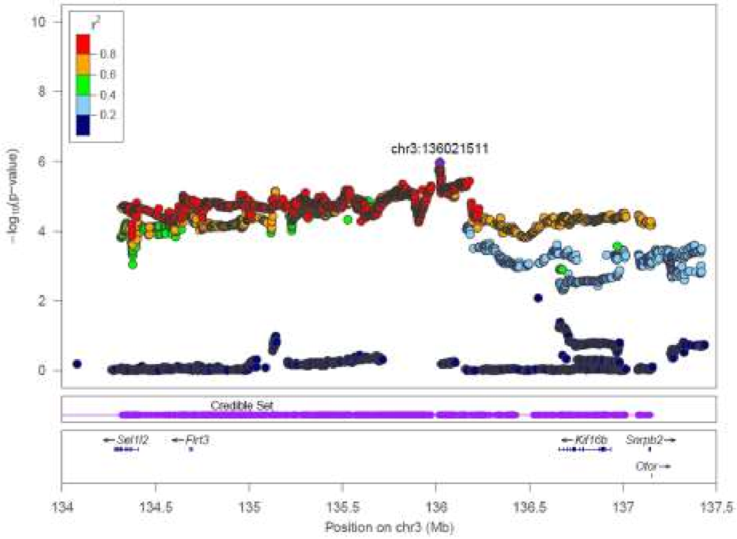


Body weight, Chr3:136,021,511

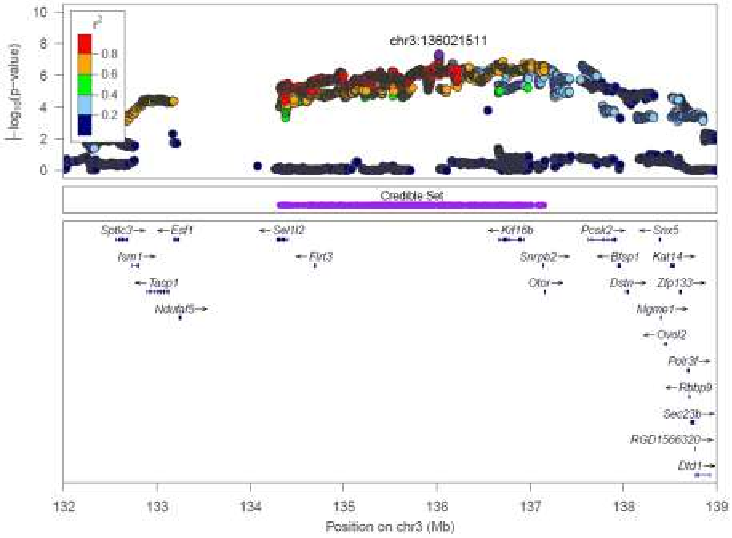

RetroFat, Chr3:137,537,161

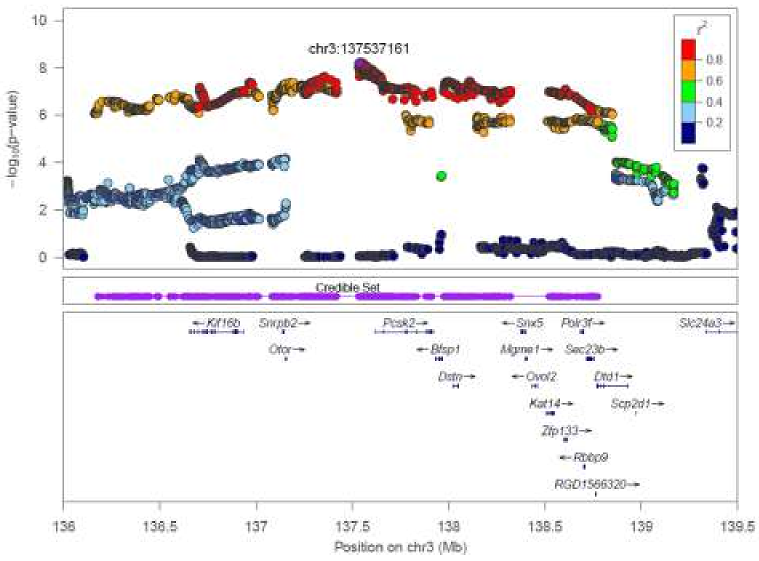


Body weight, Chr5:50,933,779

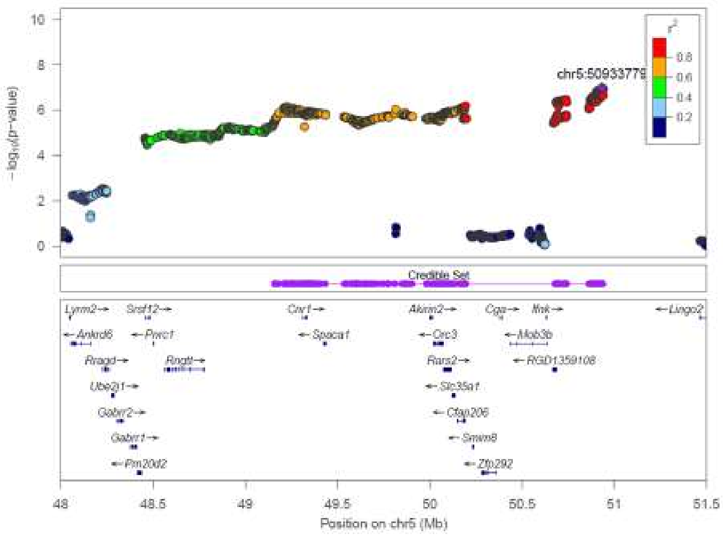

EpiFat, Chr6:26,266,960

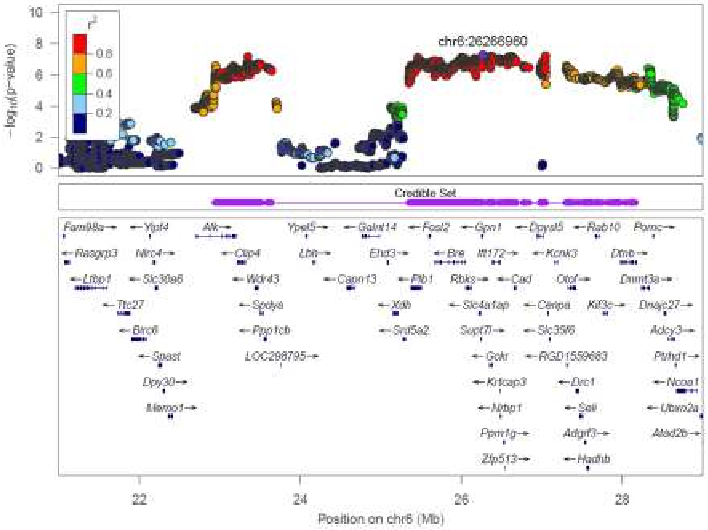

RetroFat, Chr6:28,148,338

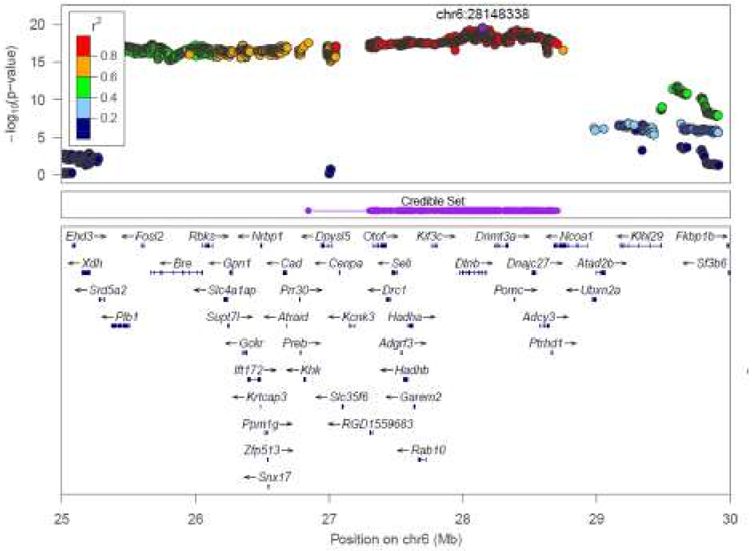

Body Length_Tail, Chr6:137,745,191

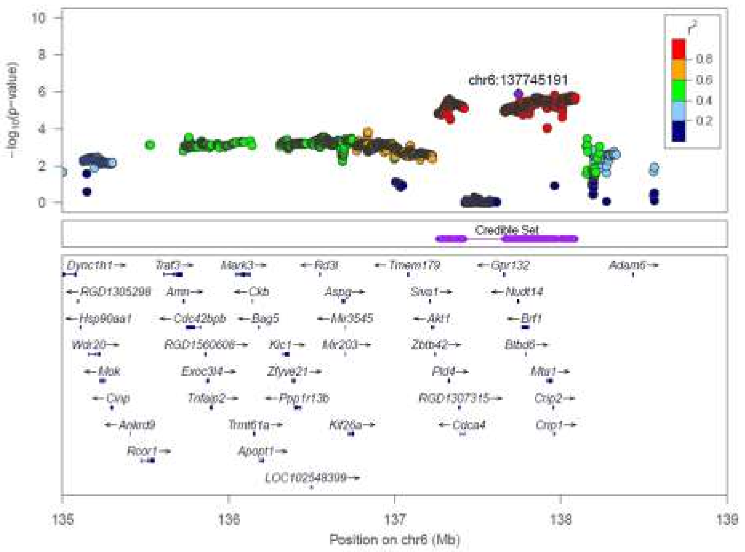


Body weight, Chr7:24,886,476

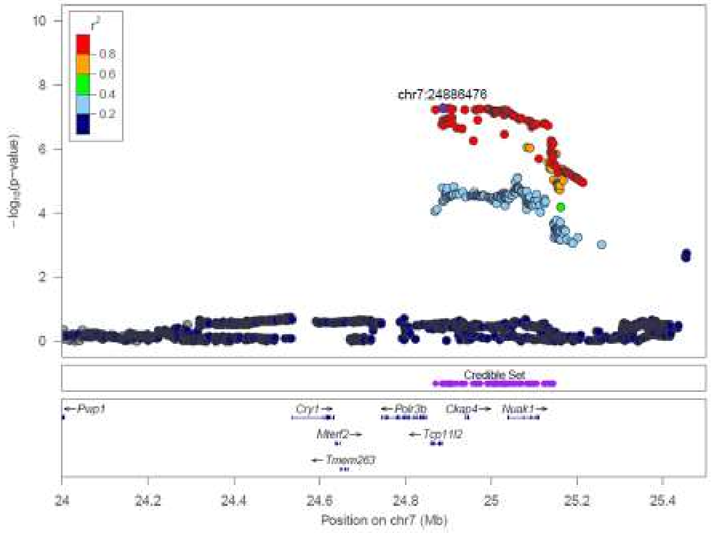

Body weight, Chr7:36,497,588

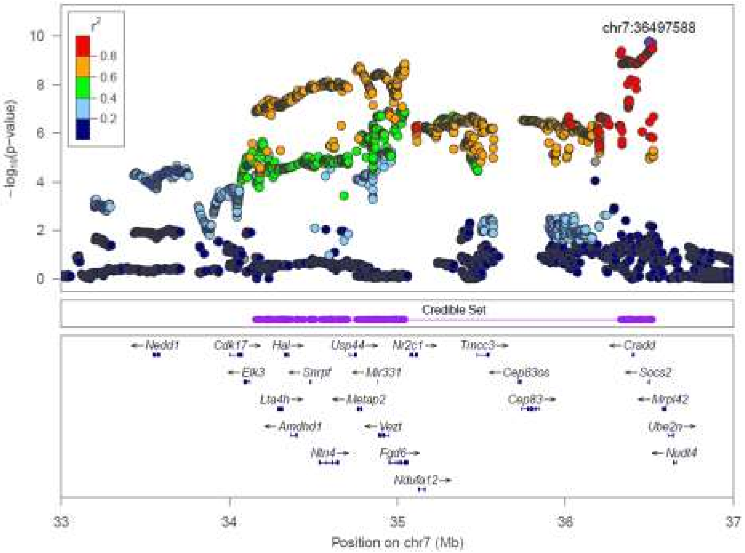


Body Length_NoTail, Chr7:36,517,726

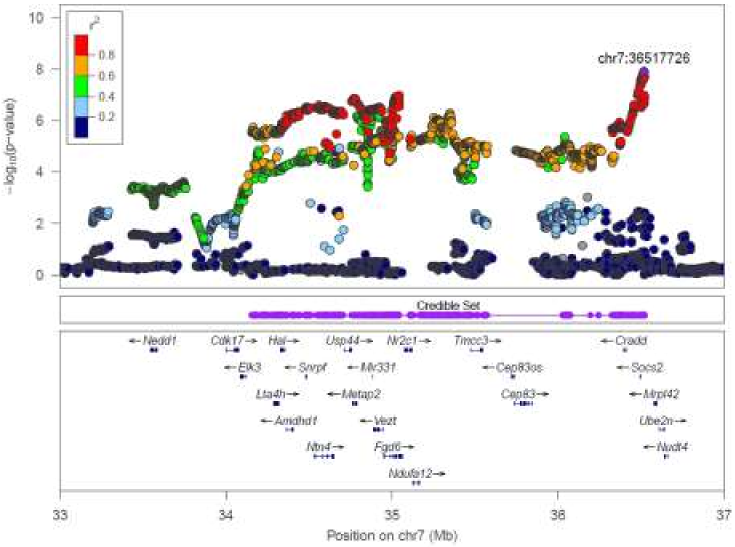

Body Length_Tail, Chr7:36,517,726

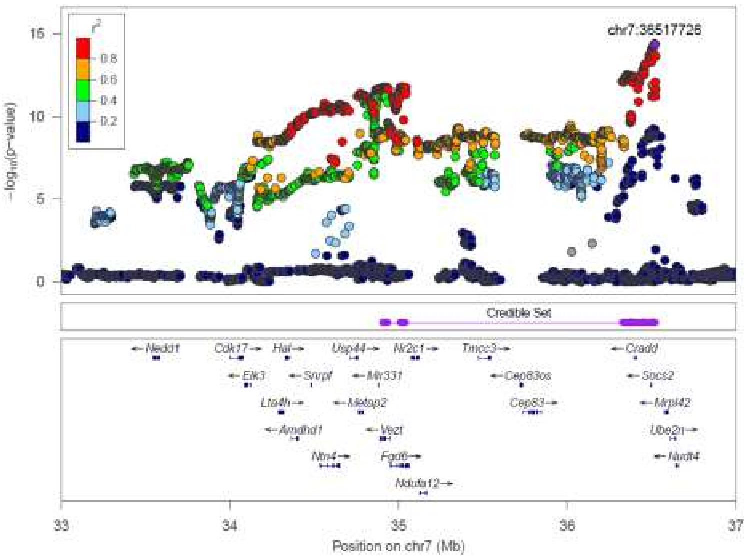


TL, Chr7:36,526,715

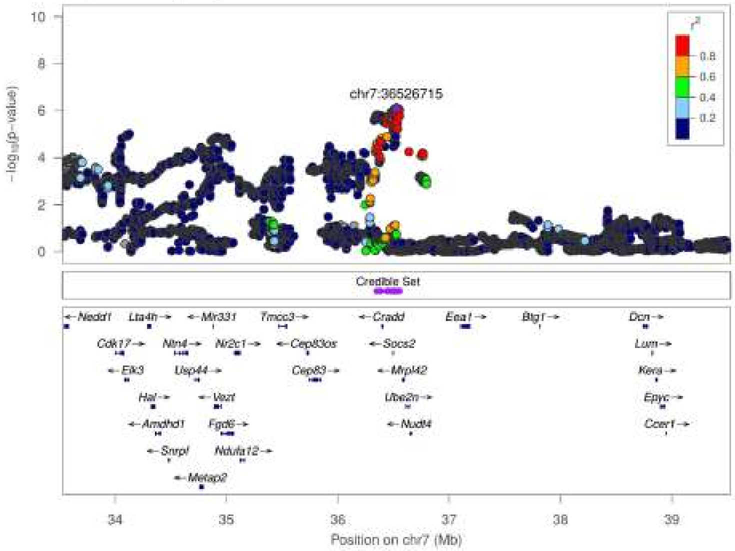

Body Length_Tail, Chr8:118,711,320

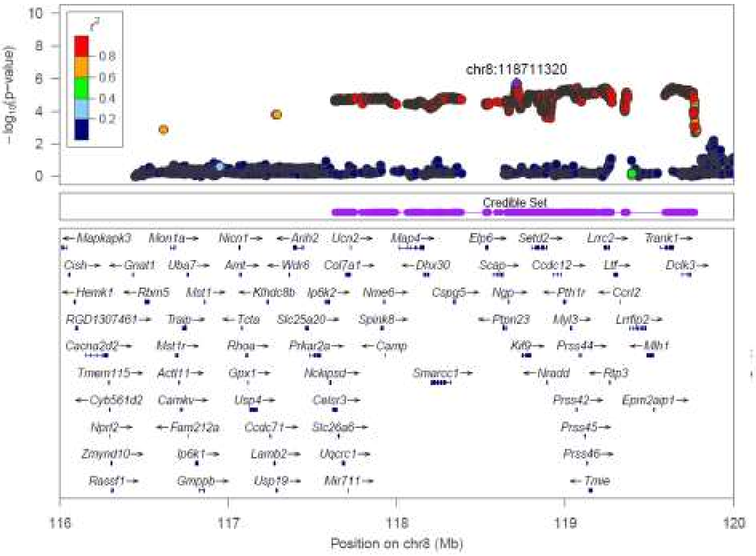


Body Length_Tail, Chr9:65,078,205

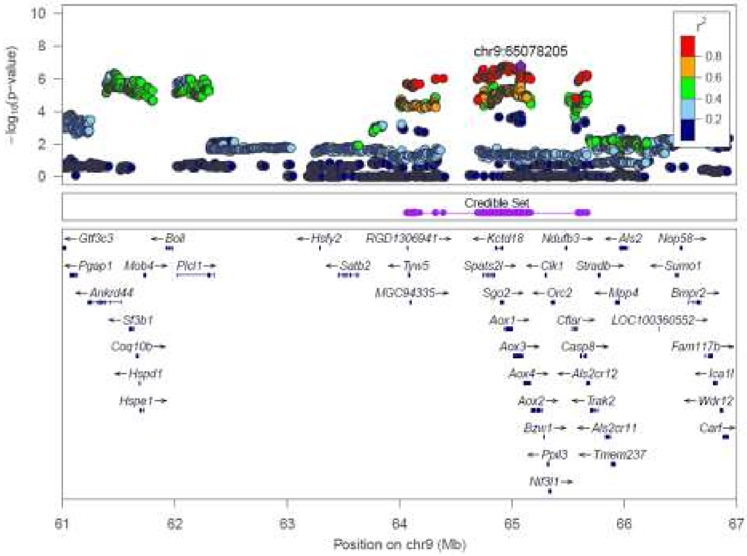

BMI_Tail, Chr10:84,080,794

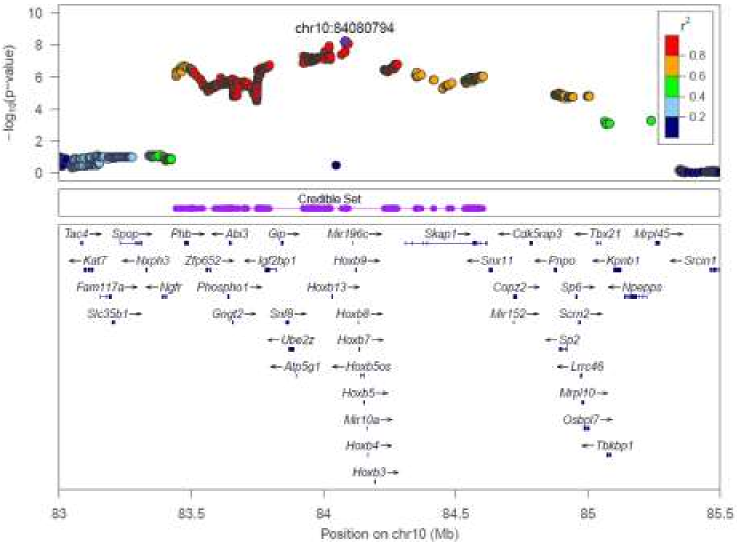


TL, Chr10:84,263,936

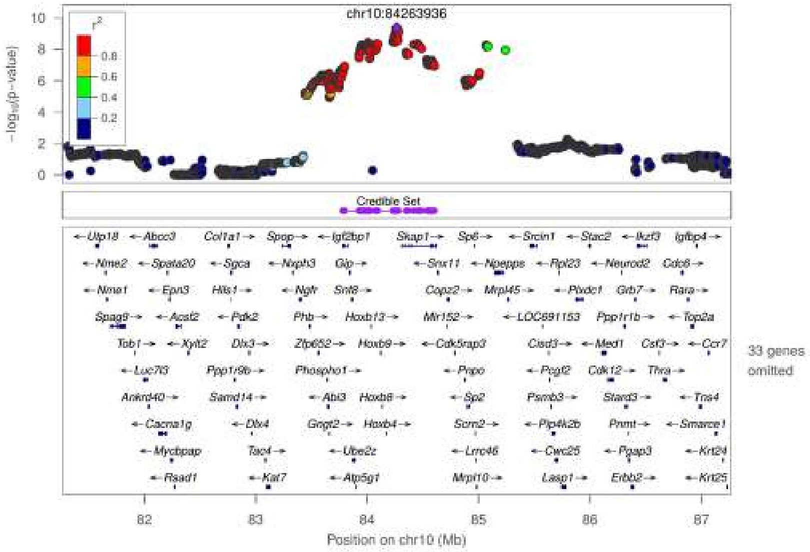

Body Length_Tail, Chr10:85,082,795

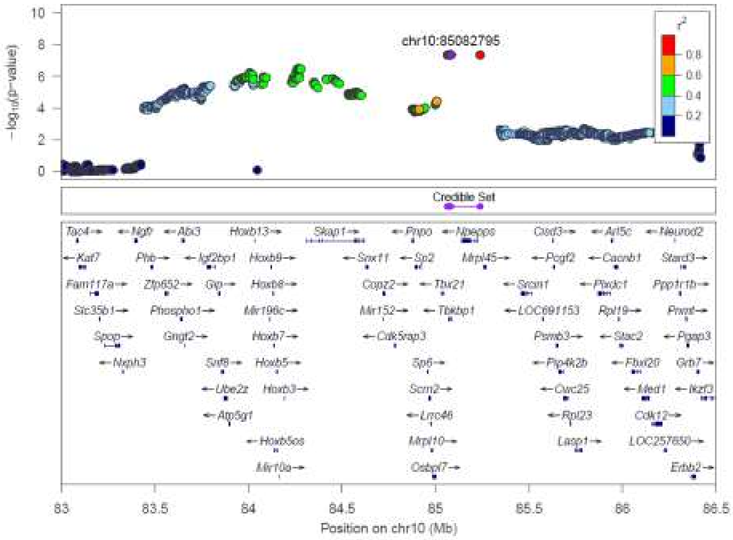


BMI_NoTail, Chr10:96,804,258

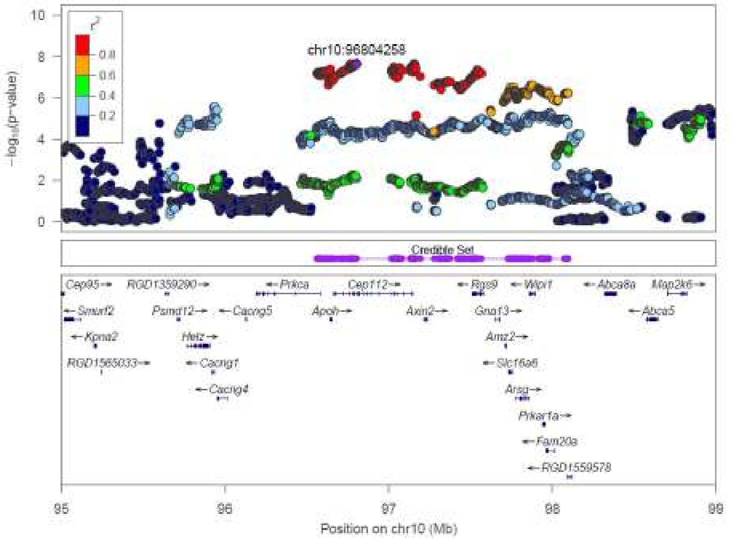

Fasting glucose, Chr10:109,944,213

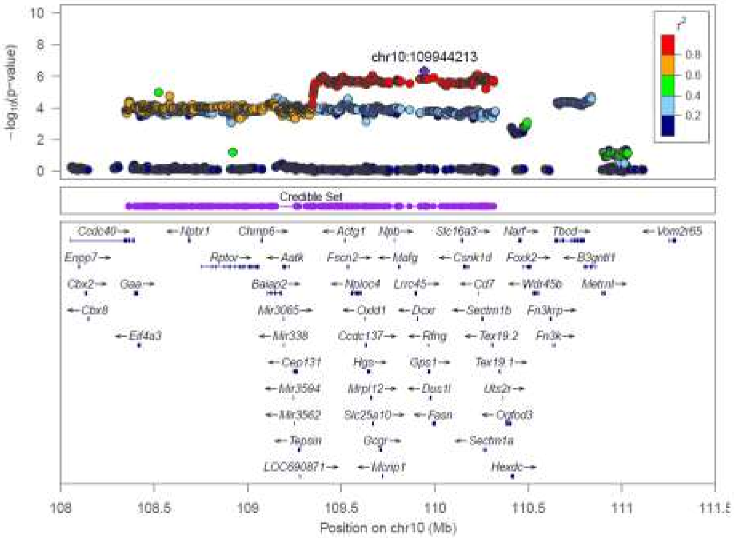


Body weight, Chr10:111,010,289

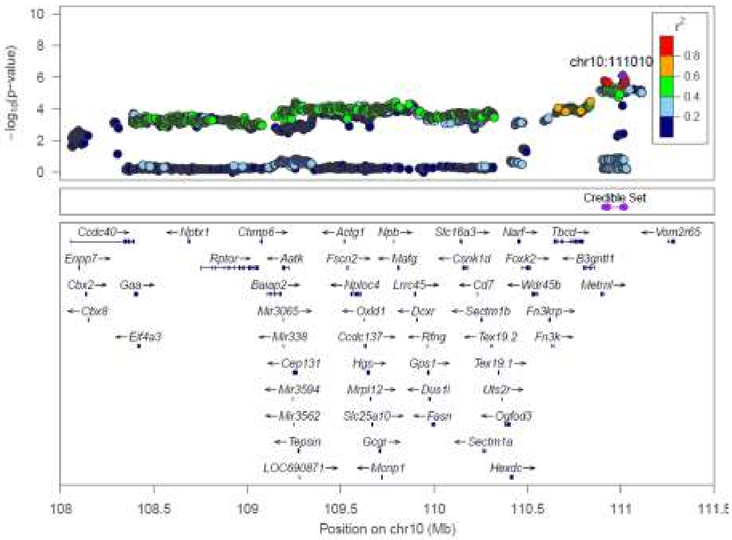

Body Length_Tail, Chr12:2,199,384

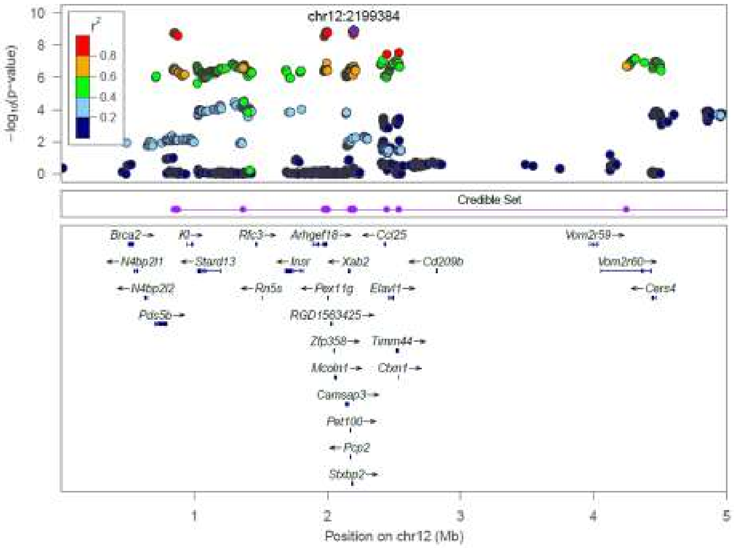


Body weight, Chr12:5,738,696

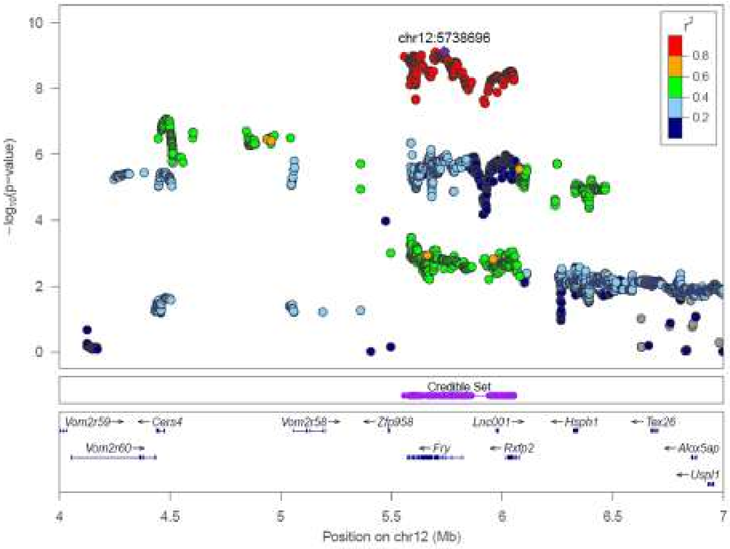

RetroFat, Chr12:5,782,829

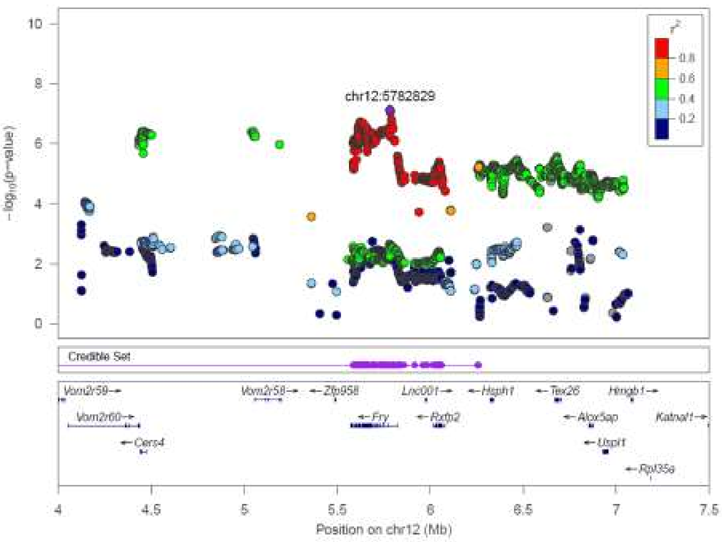


Body Length_NoTail, Chr12:6,239,515

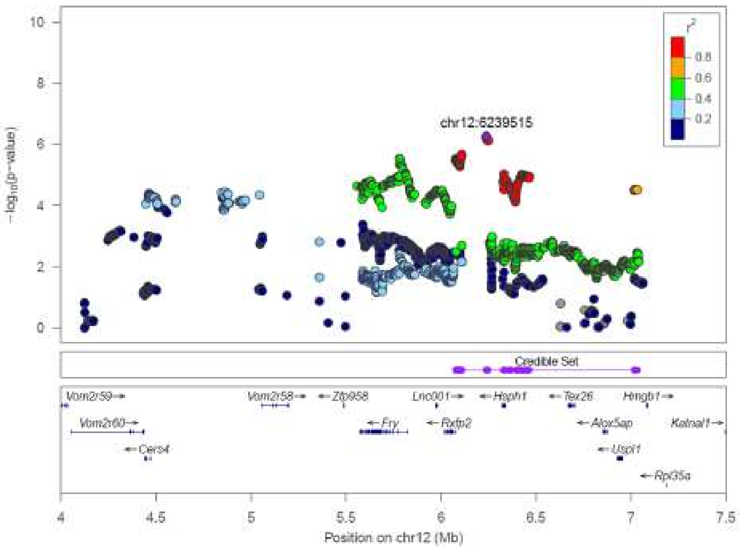

Body Length_Tail, Chr12:43,060,205

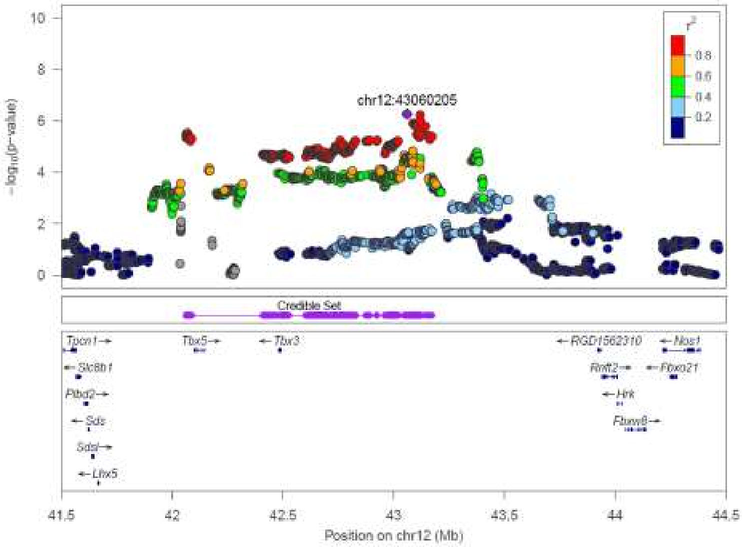


RetroFat, Chr13:55,021,887

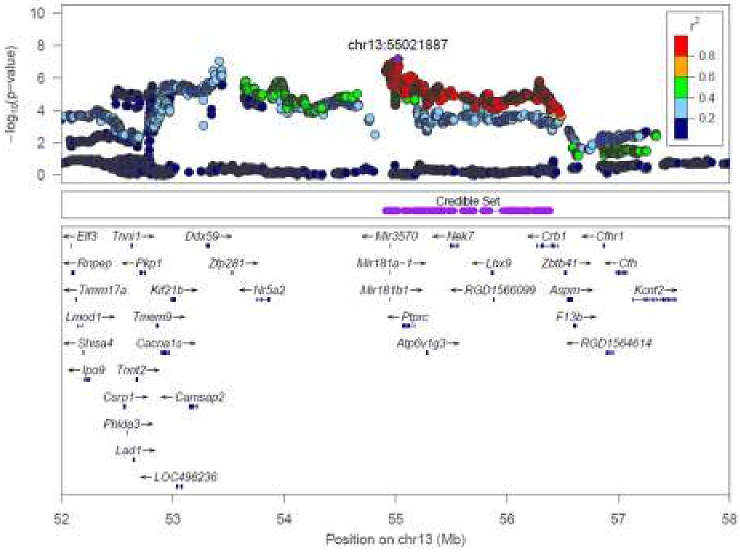

TL, Chr13:109,566,014

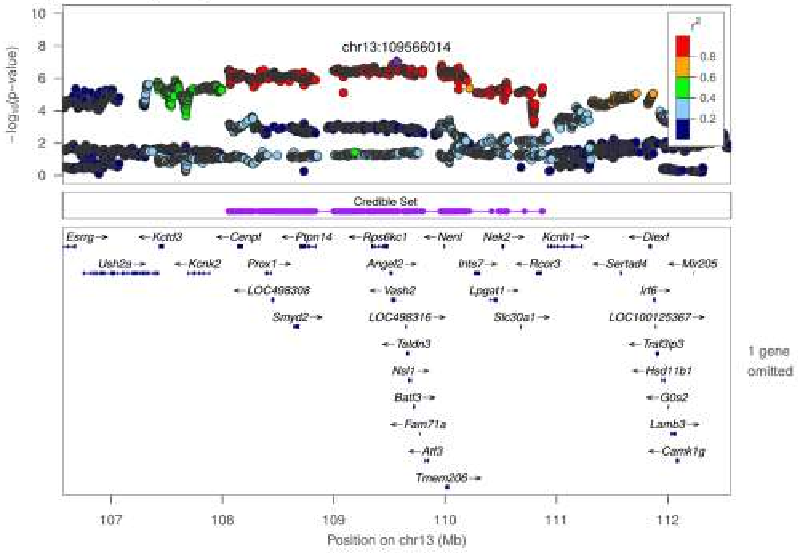


Body Length_NoTail, Chr14:26,365,986

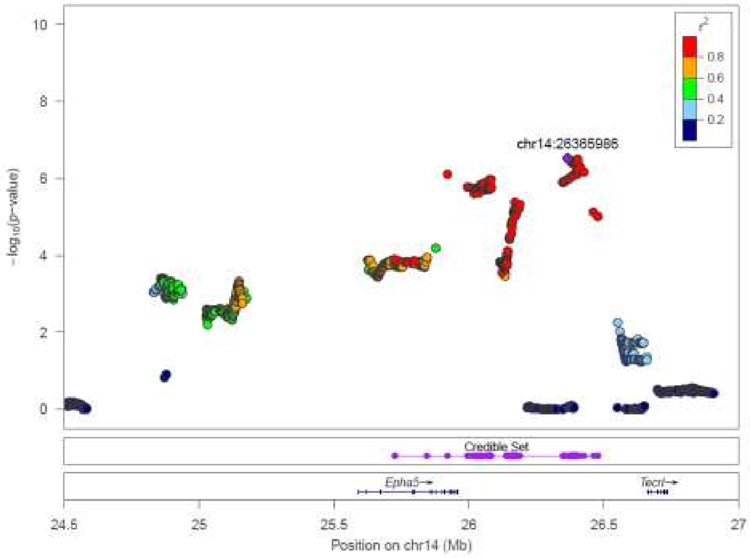

Fasting glucose, Chr14:86,029,588

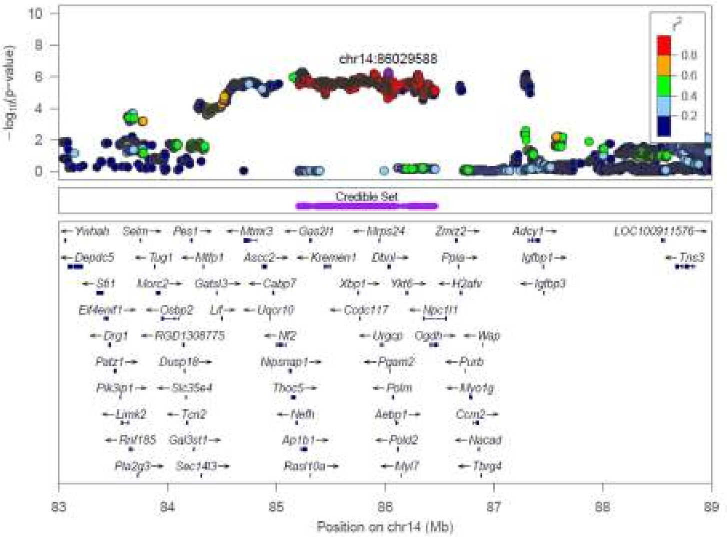

Body Length_NoTail, Chr16:64,014,119

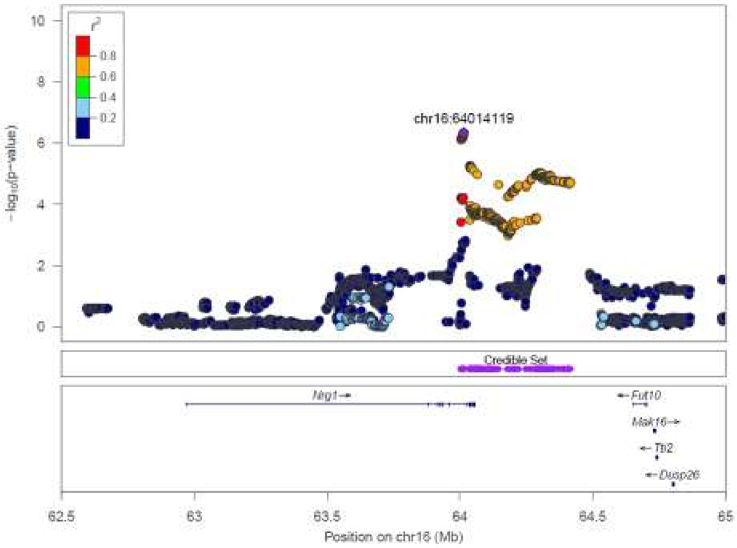

EpiFat, Chr18:12,674,871

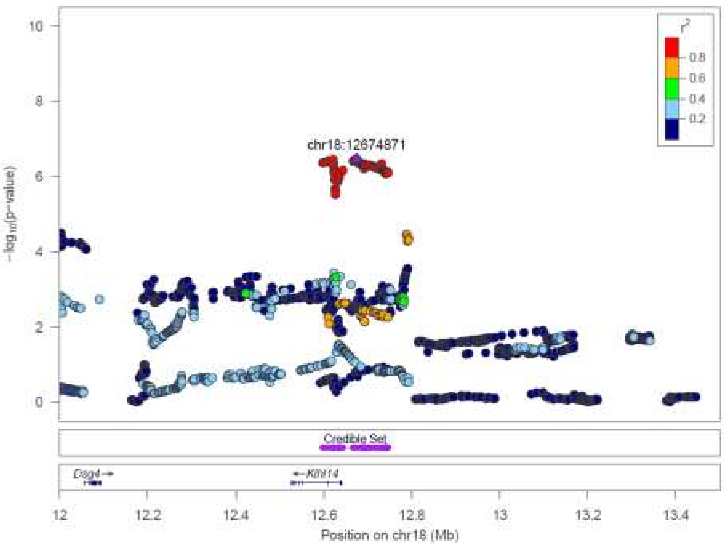


BMI_NoTail, Chr18:25,190,274

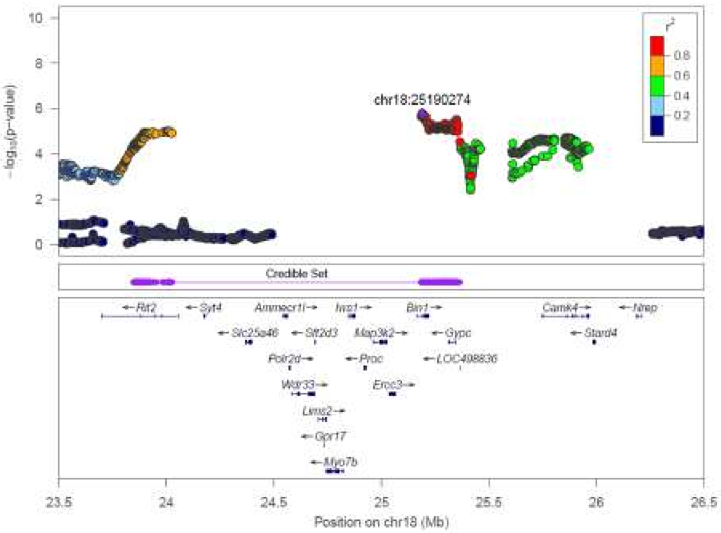

TL, Chr19:25,190,274

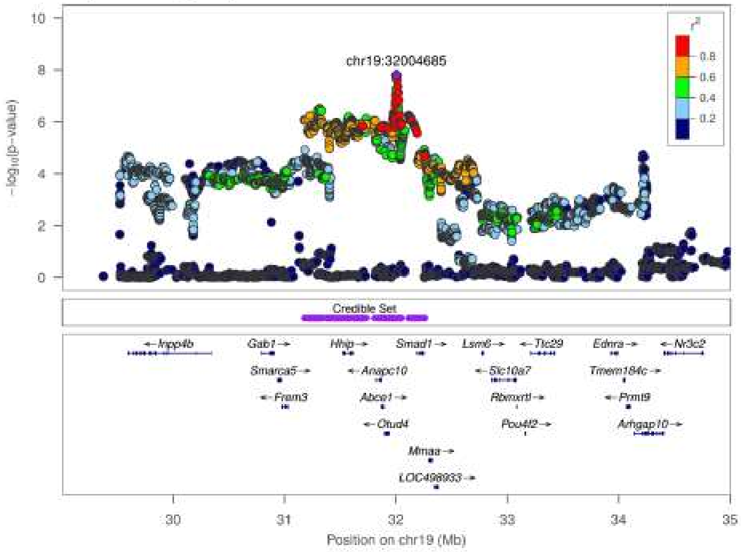
